# Dysregulated functional and metabolic response in multiple sclerosis patient macrophages correlate with a more inflammatory state, reminiscent of trained immunity

**DOI:** 10.1101/2021.01.13.426327

**Authors:** J. Fransson, C. Bachelin, F. Deknuydt, F. Ichou, L. Guillot-Noël, M. Ponnaiah, A. Gloaguen, E. Maillart, B. Stankoff, A. Tenenhaus, F. Mochel, B. Fontaine, C. Louapre, V. Zujovic

**Affiliations:** Sorbonne Université, Institut du Cerveau - Paris Brain Institute - ICM, Inserm, CNRS, APHP, Hôpital Pitié Salpétrière Univ. Hosp., DMU Neuroscience 6, Paris, France; Inst. of Cardiometabolism and Nutrition, Sorbonne-universités-Upmc 06, INSERM, CNRS, Paris, France; Laboratoire des Signaux et Systèmes (L2S), CNRS- CentraleSupélec, Université Paris-Saclay; Sorbonne Université, Institut du Cerveau - Paris Brain Institute - ICM, Inserm, CNRS, APHP, Hôpital St. Antoine-HUEP, Paris, France; INSERM, SU, AP-HP, Centre de recherche en Myologie-UMR974 and Service of Neuro-Myology, Institute of Myology, University hospital Pitié-Salpêtriere

**Keywords:** multiple sclerosis, macrophages, pro-inflammatory, pro-regenerative, homeostatic, metabolism, apoptosis, transcriptomics, trained immunity

## Abstract

In multiple sclerosis (MS), immune cells invade the central nervous system and destroy myelin. Macrophages contribute to demyelination and myelin repair, and their role in each process depends on their ability to acquire specific phenotypes in response to external signals. Here, we assess whether defects in MS patient macrophage responses may lead to increased inflammation or lack of neuro-regenerative effects.

To test this hypothesis, CD14^+^CD16^-^ monocytes from MS patients and healthy controls were activated *in vitro* to obtain homeostatic-like, pro-inflammatory and pro-regenerative macrophages. Myelin phagocytic capacity and surface molecule expression of CD14, CD16 and HLA-DR were evaluated with flow cytometry. In parallel, macrophages were assessed through RNA sequencing and metabolomics.

We observed that MS patient monocytes *ex vivo* recapitulate their preferential activation toward a CD16^+^ phenotype, a subset of pro-inflammatory cells present in MS lesions. Even in the absence of pro-inflammatory stimuli, MS patient macrophages exhibit a pro-inflammatory transcriptomic profile with higher levels of cytokine/chemokine suggesting increased recruitment capacities. Interestingly, MS patient macrophages exhibit a specific metabolic signature with a mitochondrial energy metabolism blockage resulting in a shift from oxidative phosphorylation to glycolysis. Furthermore, we observe a failure to up-regulate apoptosis effector genes in the pro inflammatory state suggesting a longer-lived pro-inflammatory macrophage population.

Our results highlight an intrinsic defect of MS patient macrophages that provide evidence of innate immune cell memory in MS.

## Introduction

Multiple sclerosis (MS) is an inflammatory disease in which peripheral immune cells infiltrate the central nervous system (CNS) and destroy myelin, a neuroprotective and conduction-enhancing substance. This leads to impaired neuronal function and finally neurodegeneration. The infiltrating cells primarily consist of lymphocytes and monocytes, which together with microglia (the tissue-resident macrophages of the CNS) induce and maintain neuroinflammation. Infiltrating monocytes differentiate into macrophages, and the total macrophage population is believed to play an important role in the disease [70]. They likely contribute to myelin destruction through perpetuation of the inflammatory environment, recruitment of leukocytes, antigen presentation and damage to neural cells through toxic effector mechanisms [4, 63].

However, demyelinated lesions can be repaired through an endogenous processed termed remyelination. The remyelination capacity varies greatly between patients, with extensive remyelination correlating with higher age at death [51] and lower disease severity [10]. In animal models, immune cells are instrumental for efficient remyelination. Depletion of macrophages leads to reduced proliferation and differentiation of oligodendrocyte precursor cells (OPCs), resulting in fewer myelin-producing oligodendrocytes [35, 46]. This effect is linked to factors secreted by macrophages and the differentiation-inhibiting properties of myelin debris [34], which is cleared through phagocytosis by macrophages in demyelinated lesions.

In this process, the activation status of macrophages appears critical. Macrophages respond to cues in the environment, such as pathogen-derived molecules and cytokines from other immune cells, and respond accordingly to achieve different “activation states” [47]. The activation states are typically described with terms such as “pro-inflammatory” (typically induced with interferon (IFN) γ and lipopolysaccharide (LPS) *in vitro*) and “anti-inflammatory” (typically induced with interleukin (IL)-4 *in vitro*). While macrophages in an organism often display features of different activation states at once, *in vitro* activation of macrophages with specific pro- or anti-inflammatory stimuli has provided great insight into the extensive macrophage response to inflammatory molecules, including the signaling pathways that induce this functional diversity [21, 44, 54].

Pro-inflammatory macrophages produce more pro-inflammatory cytokines and toxic molecules, suggesting a destructive role in MS. In the case of effects on remyelination, *in vivo* and *in vitro* results show that pro-inflammatory macrophages promote OPC proliferation while anti-inflammatory macrophages promote OPC differentiation. *In vivo* data show a switch from a majority of pro-inflammatory cells to a population composed of pro-regenerative macrophage during the early stages of successful remyelination [46]. This switch depends on a necroptotic depletion of pro-inflammatory macrophages, permitting the resolution of inflammation and establishment of an anti-inflammatory population [42]. Thus, pathological activation of macrophages appears as a potential culprit in both destruction and lack of repair.

Macrophage activation is indeed extensive in MS CNS, and loss of homeostatic marker purinergic Receptor P2Y12 (P2RY12) is seen both in lesions and in normal-appearing white matter [74]. Pro-inflammatory macrophage markers are abundant in active lesions and slowly expanding lesion rims [26, 74], both of which show active demyelination. However, it is not known to what extent this perturbed state is driven by a pathological environment or by intrinsic features of infiltrating macrophages.

Recently, it was shown that innate immune cells (monocytes, macrophages and microglia) can develop an immunologic memory named trained immunity [50]. This phenomenon has been observed in inflammatory disease such as atherosclerosis [8]. Induction of this trained state relies on epigenetic reprogramming resulting in a metabolic switch toward glycolysis and an increased activation of innate immune cells characterized by an enhanced inflammatory response and cytokine production. However, the differences in macrophage capacity for homeostasis and activation have not been studied in MS conditions, and thus it is unknown whether monocyte-derived macrophages are primed to react incorrectly.

In this study, we examine whether MS patient macrophages show differences compared to healthy controls (HC), and whether these are dependent on activating stimuli. By using monocytes isolated from blood samples and differentiating them *in vitro*, we emulate infiltrating macrophages without exposure to the lesion. Our original approach explores how disease relates to key components of macrophage activation at the functional and molecular levels and how these dysfunctionalities might lead to the onset or development of proinflammatory lesions. We provide evidence of a pro-inflammatory phenotype in MS patient macrophages, with transcriptomic, metabolic and functional changes.

## Methods

### Participants

A total of 36 MS patients and 17 HC were included in this study (Supplementary table 1). The study was approved by the French Ethics committee and the French ministry of research (NCT03369106). Written informed consent was obtained from all study participants. Patients were recruited from a larger cohort of multi-case families, collected within a project studying MS patient phenotype in multiplex families. Six of the patients were the only one in their family to be sampled, the remaining 30 were 15 pairs of siblings. All patients fulfilled diagnostic criteria for MS, and individuals (MS patients and healthy donors) with other inflammatory or neurological disorders were excluded from the study. For patients, clinical evaluation and blood sampling were performed on the same day. The clinical evaluation included standard testing of Expanded Disability Status Scale (EDSS) [37], from which MS Severity Score (MSSS) [59] was calculated, as well as documentation of treatment history. The varying number of patients and control in the different experiments are mainly due to cell number constraint or experimental limitations.

**Table 1.** Characteristics of study participants. Continuous variables are given as mean ± standard deviation.

|  | Age<br>(years) | Female<br>sex | Disease<br>Duration<br>(years) | % MS-treated | % RRMS | MSSS |
| --- | --- | --- | --- | --- | --- | --- |
| HC | 37±11 | 14/17<br>(82%) | | | | |
| MS | 38±11 | 31/36<br>(86%) | 12±8 | 25/36 (69%) | 33/36 (92%,<br>2 CIS 1 SP) | 2,9±1,7 |

*RRMS: Relapsing remitting; CIS: Clinically isolated syndrome; SP: secondary progressive* *MSSS: Multiple sclerosis severity score*

### Macrophage culture and activation

Blood was sampled from all participants in acid citrate dextrose (ACD) tubes. From blood samples, peripheral blood mononuclear cells (PBMCs) were isolated using Ficoll Paque Plus (GE Healthcare Life Sciences) and centrifugation (2200 rpm, 20 min). Cells were washed in PBS (2×10 min at 1500 rpm) and RPMI 1640 + 10% fetal bovine serum (FBS) (5 min at 1500 rpm) (all products from ThermoFisher). Monocytes were isolated with anti-CD14 microbeads (Miltenyi) and plated in 12-well plates (500 000 cells/well) or in 24-well plates (200 000 cells/well) in RPMI 1640 + 10% FBS and granulocyte macrophage colony-stimulating factor (GM-CSF) (500 U/ml, ImmunoTools). After 72h, media was replaced with fresh media and one of the following: GM-CSF (500 U/ml); IL4 (1000 U/ml, ImmunoTools); or combined IFNγ (200 U/ml, ImmunoTools) and ultra-pure LPS (10 ng/ml, InvivoGen). Purity after isolation was evaluated by FACS analysis with anti-CD14-FITC (Miltenyi), anti-CD16-eFluor450 (ThermoFisher) and anti-CD3-APC (BioLegend).

### Human myelin extraction

Human myelin was extracted from normal-appearing white matter in post mortem MS patient brain tissue. Tissue was homogenized in 0.32M sucrose (prepared in 20mM Tris-HCl). Homogenate was layered on 0.85M sucrose (prepared in 20mM Tris-HCl) and sample was centrifuged at 75,000 g for 30 min. The layer between the two sucrose layers was collected and washed with water 3 times, first at 75,000 g and then at 12,000 g twice. Pellet was resuspended in 0.85M sucrose and 0.32M was layered on the solution. Sample was centrifuged at 75,000 g for 30 min. Pellet was resuspended in water and centrifuged at 75,000 g for 15 min. Total myelin protein was quantified with Micro BCA™ Protein Assay Kit (ThermoFisher Scientific). Myelin was labeled using Amersham CyDye Cy5 Mono-Reactive Dye (GE Healthcare Life Sciences), using half of the recommended dose (1 vial per 2mg myelin). Labeled myelin was washed with PBS and resuspended in PBS at 1µg/µl.

### Phagocytosis assay and flow cytometry

Macrophage phagocytic capacity was evaluated through flow cytometry. Twenty-four hours post-activation, media was replaced for macrophages in 24-well plates with RPMI + 10% FBS containing labeled human myelin (25 µg/ml). For each condition, 1 well was used for myelin incubation and one well was used for negative control. Cells were incubated at 37C for 1 hour. Cells were washed with PBS and detached from wells using Trypsin-EDTA (0.25%) (ThermoFisher). Cells were flushed with RPMI 1640 + 10% FBS until all cells had detached, at which point cells were washed through centrifugation with PBS. Labeling was done with anti-HLA-DR-PB and anti-CD16-FITC (Duraclone pre-coated tubes or separate antibodies, all from Beckman-Coulter) and anti-CD14-PEvio770 (Miltenyi) at room temperature in darkness for 30 min (pre-coated tubes) or 45 min at 4°C (liquid antibodies) in PBS + 5% FBS. Cells were washed with PBS + 5% FBS and analyzed with MACSQuant (Miltenyi). Results were analyzed with Flowlogic software (Inivai Technologies). Cells were gated for size and granulosity (FSC/SCC) and singlets (FSC-A/FSC-H).

### Multiway Generalized Canonical Correlation Analysis

Flow cytometric data was analyzed using Multiway Regularized Generalized Canonical Correlation Analysis (MGCCA) [22]. Prior to analysis, samples with low cell counts or missing data points were excluded, and 33 patients and 16 HC were used in the final analysis. Data were log2-transformed and formatted into a 3-order tensor with dimension 1 corresponding to individual participants, dimension 2 to flow cytometric variables and dimension 3 to activation states. One of the main goals in the method is to understand the complex relationships between this flow cytometric 3-order tensor data and the response variable (HC vs MS).

Regularized generalized canonical correlation analysis (RGCCA) is a general framework for multiblock data analysis [65, 66]. RGCCA is geared for the analysis of a set of matrices.

RGCCA can also be applied to tensor data using the multiway formalism, which accounts for multiple measurements of the cytometric variables along activation states. This method, called MGCCA, was used to explore the complex relationships between flow cytometric data measured at 3 different states of activation with the response variable (HC vs MS). MGCCA allows identifying (through the construction of two weight vectors associated with cytometric variables and activation states) cytometric variables within specific activations state that are discriminant between HC and MS.

A bootstrap procedure [16, 17] was performed to assess the reliability of parameter estimates. 500 bootstrap samples of the same size as the original data were repeatedly sampled with replacement from the original data. MGCCA was applied to each bootstrap sample to obtain estimates (i.e. cytometric and activation states weights). We then calculated the mean and variance of the estimates across the bootstrap samples, from which we derived confidence interval. A coefficient is declared significant when zero is excluded from its confidence interval.

### RNA Sequencing and differential expression analysis

Cell lysis and RNA extraction were performed on macrophage samples 24h post-activation using Nucleospin RNA extraction kit (Macherey-Nagel). Quality of RNA was confirmed on Agilent TapeStation (RINe>8). Transcriptome sequencing was performed on a total of 28 patients and 11 HC. cDNA libraries were prepared using a stranded mRNA polyA selection (Truseq stranded mRNA kit, Illumina). For each sample, we performed 60 million single-end, 75 base reads on a NextSeq 500 sequencer (Illumina).

Quality of raw data was evaluated with FastQC. Poor quality sequences were trimmed or removed with Fastp software to retain only good quality paired reads. Star v2.5.3a [13] was used to align reads on reference genome hg38 using standard options. Quantification of gene and isoform abundances has been done with RSEM 1.2.28 [41].

In the analyses detailed below, two batch effects were taken into account: 1) Sequencing was performed twice, with 18 RNA samples sequenced in both rounds; 2) In the second sequencing round, cDNA libraries were prepared in two batches, with one batch showing a noticeable difference in DNA quantity. In addition, one patient was sampled twice – the first sample was sequenced in the first run, and the patient was later resampled in order to increase the number of untreated patients, as (independently of the study) the patient had suspended treatment between the two visits. The data from first sample were therefore excluded in the comparisons described below.

Differential expression analysis was performed using limma (3.40.6) [57] in combination with voom [40], including the batch effects in the design and improving batch correction using the DuplicateCorrelation function with the re-sequenced samples. Only genes expressed at a minimum of 1 CPM were included in the analysis. Multiple hypotheses-adjusted p-values were calculated with the Benjamini-Hochberg procedure to control FDR. Results were considered statistically significant at p-value <= 0.05 and Log2FC >=0.5.

Functional enrichment analysis was performed with EnrichR [36]. GO Biological Process terms were considered significantly over-represented at adjusted p-value <0.05. Terms were simplified with the simplifyGOterms function from the compEpiTools package.

When two analyses were compared (e.g. M_IL-4_ vs M_GM-CSF_ in MS and M_IL-4_ vs M_GM-CSF_ in HC), genes were considered specific to MS if the fold-change of significant genes in MS was 0.5 higher (for over-expressed genes) or 0.5 lower (for under-expressed genes) than in HC (and vice versa for HC).

To illustrate gene expression while accounting for batch effects, a representative data set was produced using the removeBatchEffect function of the limma package with the same parameters as lmFit.

### Principal component analysis

To produce a batch-corrected transcriptomic dataset without bias toward the variables of interest, the removeBatchEffect function from the limma package was again used, correcting for the batch effects described above, and including the intra-duplicate correlation. Unlike the data set used to visualize specific genes (above), the patient status and activation state were not considered in the correction.

Principal component analysis (PCA) was performed in R [56] using the ExPosition R package [7]. All analyses were performed using centered but not scaled log_2_-transformed RPKM+1 values. Only genes that were expressed in ≥80% of all samples were included. Samples from one individual were either considered individually or combined into one observation by defining each gene in each activation state as a separate variable. To calculate 95% confidence intervals for each group mean, a bootstrap procedure [16, 17] was performed. 100 bootstrap samples of the same size as the original data were repeatedly sampled with replacement from the original data. PCA was applied to each bootstrap sample and the group means were recalculated from these samples, from which we derived confidence interval.

### Weighted gene co-expression network analysis

Weighted gene co-expression network analysis (WGCNA) [38] was performed. We used the same batch-corrected data as for the PCA. In order to reduce the effect of the activation state in the network, the expression data was divided into three sets, one per activation state, and a multi-dataset network was constructed in a similar way to the consensus network instructions from the WGCNA package creators [39]. However, instead of considering the minimal correlation across datasets for each gene pair, the maximal correlation was considered. Genes were filtered to exclude any gene not expressed in ≥80% of samples in at least one dataset. Functional annotation of genes in WGCNA modules was performed using EnrichR to identify over-represented KEGG pathways [28] and GO biological process and molecular function terms [6]. Terms were considered significantly over-represented at FDR-adjusted p<0.05.

### Metabolomics

Metabolomics analysis was performed on GM-CSF-exposed macrophages from 7 patients and 6 HC. Macrophage samples were prepared in the same way and in parallel to RNASeq samples until the lysis step. At this point, cells were instead detached from wells with 0.25% Trypsin-EDTA (ThermoFisher) for 10 min in 37°C. Cells were collected in RPMI + 10% FBS and centrifuged at 400g to form a pellet. After removal of supernatant, samples were stored at - 80°C. The numbers of cells for each sample were estimated based on the number of cells measured in the corresponding flow cytometry sample, adjusted for the number of cells plated for each experiment.

Prior to extraction process, dry pellet cells were homogenized in 0.1% formic acid including internal standards labelled mix of amino acids (10 µg/mL) to a ratio equivalent to 50.000 cells per 100µL. 4 volumes of frozen methanol (-20°C) containing internal standards as well (labeled mixture of amino acids at 10 µg/mL) were added to 100 µL cell samples and vortexed. The resulting samples were sonicated during 10 min and centrifuged during 2 minutes at 10.000xg and at 4°C. Then, centrifuged samples were incubated at 4°C during 1 hour for the slow protein precipitation process. Samples were centrifuged for 20 min at 20.000 × g at 4°C. Supernatants were transferred to another series of tubes and then dried and stored at -80°C prior to the LC-MS analyses. Aliquots were reconstituted in 200µL of H2O/ACN (40/60).

Liquid chromatography mass spectrometry (LC-MS) experiments were performed using a HILIC phase chromatographic column, on a ZIC-pHILIC 5µm, 2.1 × 150 mm at 15°C (Merck, Darmstadt, Germany), and on a UPLC Waters Acquity (Waters Corp, Saint-Quentin-en-Yvelines, France) coupled to Q-Exactive mass spectrometer (Thermo Scientific, San Jose, CA). Experimental settings for global approach by LC-HRMS were carried out as detailed in the paper of Garali et al [19].

All LC-MS grade reference compounds, water (H2O) and methanol (MeOH) were from VWR International (Plainview, NY). LC grade 2-propanol (IPA) and formic acid were from Sigma-Aldrich (Saint Quentin Fallavier, France). Stock solutions of Stable isotope-labeled mix (Algal amino acid mixture-^13^C,^15^N) for metabolomics approach were purchased from Sigma-Aldrich (Saint Quentin Fallavier, France).

All processing steps were carried out using the R software [56]. LC-MS raw data were firstly converted into mzXML format using MSconvert tool [33]. Peak detection, correction, alignment and integration were processed using Workflow4metabolomics (W4M) platforms [20]. The resulted datasets were log-10 normalized, filtered and cleaned based on quality control (QC) samples as described in Dunn et al [14]. Metabolomics features were annotated based on their mass over charge ratio (*m/z*) and retention time using an “in house” database detailed in the paper of Boudag et al [11], and also characterized based solely on their m/z using public databases such as the human metabolome database HMDB [72] and the Kyoto Encyclopedia of Genes and Genomes database, KEGG [28].

### Other statistical analyses

Non-transcriptomic continuous variables were compared between two groups using Mann-Whitney U test, and were considered significantly different at p<0.05. Non-transcriptomic correlation calculations were performed with Pearson’s correlation coefficients and p-values based on linear regression (H_0_: β=0). All analyses were performed in R v 3.6.

## Results

### Generation of monocyte-derived macrophages

Monocytes were isolated from peripheral blood mononuclear cells (PBMCs) of 36 MS patients and 17 age- and sex-matched HC (Table 1 and S1) using CD14-specific magnetic separation. For 7 HC and 26 MS patients, isolated cells were analyzed for CD14, CD16 and CD3 positivity using flow cytometry in order to characterize monocyte and lymphocyte populations (gating strategies shown in Fig. S1A, B). Among these individuals, PBMCs were analyzed in the same way for 7 HC and 19 patients. No differences were seen in PBMCs pre-isolation, with similar proportions of CD3^+^ lymphocytes and CD14^+^ and/or CD16^+^ monocyte lineage cells, as well as similar distributions of CD14- and CD16-positivity among monocyte lineage cells (Fig. S1a, c). After isolation, patient samples showed a slightly reduced proportion of CD14^+^CD16^-^ cells compared to HC, but this was not explained by an increase of another specific population (Fig. S1b, d).

Once isolated, monocytes were differentiated *in vitro* through exposure to GM-CSF for 72 hours. Macrophages were then exposed to activating stimuli (IFNγ+LPS for pro-inflammatory activation or IL4 for anti-inflammatory activation) or maintained in GM-CSF for 24h. We refer to these samples as M_GM-CSF_, M_IFNγ+LPS_ and M_IL4_ according to the stimuli used.

### MS macrophages differ from HC in function and expression of cell surface markers

Having started off with a primarily CD14+CD16-monocyte population when generating the macrophages, we examined how expression of CD14 and CD16, as well as HLA-DR differed between HC and MS macrophages after each activation stimuli. We also studied cell’s capacity to phagocytose myelin, as this capacity is likely relevant to both destruction and regeneration in MS. Samples were incubated with Cy5-labeled human myelin for 1h and then labeled with antibodies targeting CD14, CD16 and HLA-DR. Samples from 33 patients and 16 HC were evaluated on percentage of phagocytic (myelin-positive) and high-phagocytic (myelin-high) cells and on percentage of CD14-negative, -low or -high and CD16-negative or -positive cells (Fig. 1a, full gating strategies shown in Fig. S2a). Mean fluorescence intensity (MFI) was also measured for each of the surface markers.

**Fig. 1.**
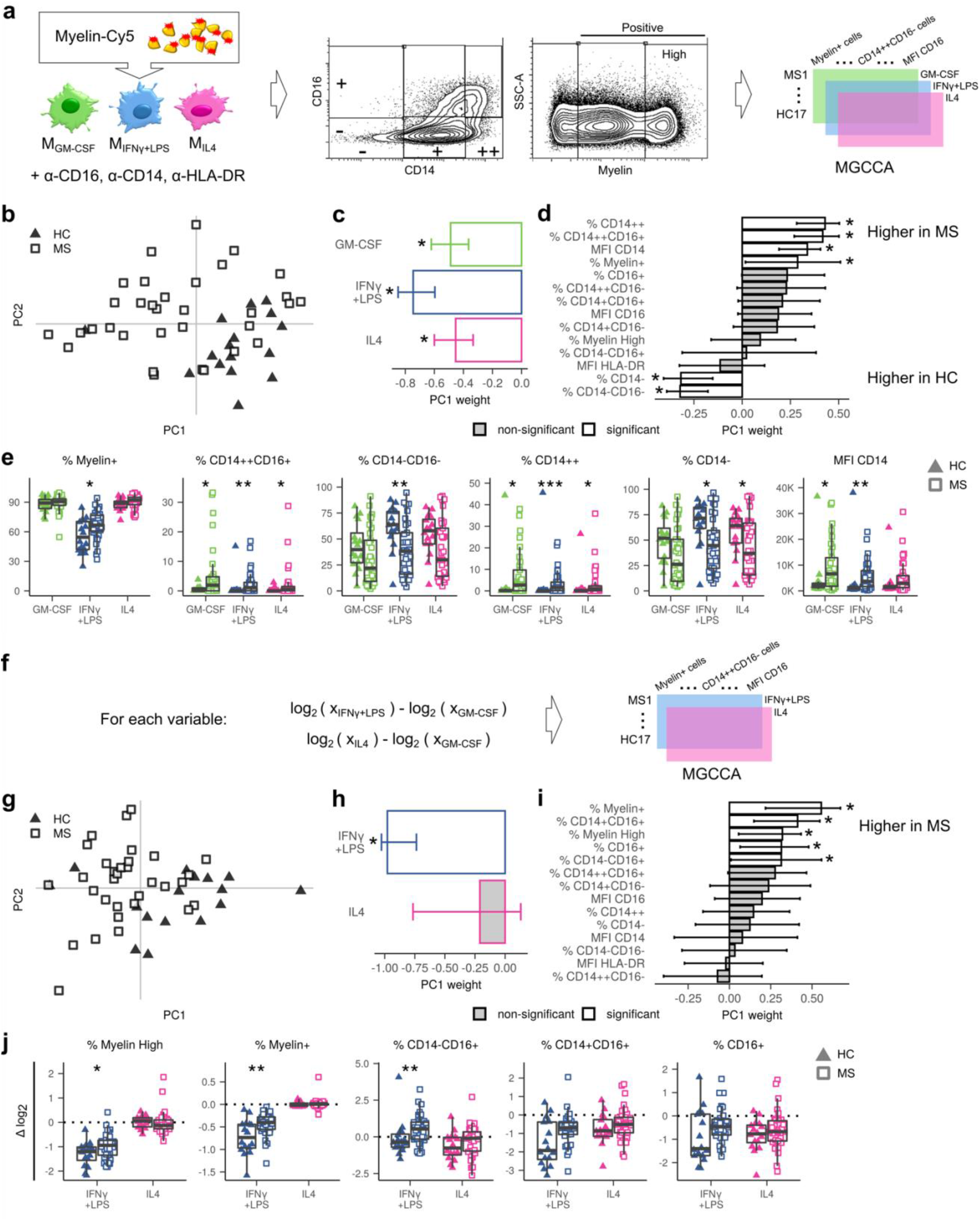
Flow cytometry reveals differences in phagocytic function and cell surface markers of HC and MS macrophages. **a)** Experimental procedure overview: Activated macrophages were exposed to labeled human myelin for 1h, and labeled with antibodies targeting CD14, CD16 and HLA-DR. In addition to registering MFI for each surface marker, the percentage of CD14 negative, low and high and CD16 negative and positive cells, and myelin positive and high, populations were calculated for each sample. A 3D matrix was constructed with dimensions representing individuals, activation states and flow cytometric variables was constructed to perform MGCCA. **b)** First principal components of MGCCA targeting the differences between MS (squares) and HC (triangles) across all activation states and variables. **c)** Weights of each activation state to the first PC of b. **d)** Weights of each flow cytometry variable to the first PC of b. **e)** Distribution of samples for each significant variable, as identified in d. Values are grouped by activation state and disease (HC: triangles; MS: squares; GM-CSF: green; IFNγ+LPS: blue; IL4: pink). **f)** Analysis procedure for inter-state differences: Data from A were transformed by subtracting the values in the GM-CSF state from the values of the same individual in the IFNγ+LPS and IL4 states. A matrix of the relative data was created as in A. **g)** First principal components of MGCCA based on relative data. **h)** Weights of each activation state to the first PC of G. Error bars indicate 95% confidence interval as calculated by bootstrapping. **i)** Weights of each flow cytometry variable to the first PC of G. **j)** Distribution of samples for each significant variable, as identified in i. Values are expressed as log2(fold-change from M_GM-CSF_) and are grouped by activation state and disease as in e. *0 outside of 95% confidence interval based on bootstrapping (d&i) or * p<0.05, ** p<0.01 in Mann Whitney U test between HC and MS for each activation state, not adjusted for multiple testing (e&j). HC: healthy controls; MS: multiple sclerosis patients. (HC n=16; MS n=33). Error bars indicate 95% confidence interval as calculated by bootstrapping.

In order to efficiently identify the key differences among these multiple parameters, multiway regularized generalized canonical correlation analysis (MGCCA) was performed with disease status (HC or MS) as the target feature. In this analysis, the three activation states were considered for each individual by constructing a 3-order tensor (individuals×biological markers×activation states) (Fig. 1a, right panel). This method generates principal components permitting visualization of the individuals in a low dimensional space. In the space of the first component, all but one HC (Fig. 1b, black triangles) clustered on one end of the scale. The patients (Fig. 1b, white squares) were more heterogeneous but on average presented on the opposite end of the scale. The 3 activation states all contributed to the separation between HC and MS patients (Fig. 1c). Looking into all the markers, we identified that MS patient macrophages were defined by a high percentage of CD14^++^ and CD14^++^/CD16^+^ populations, a high MFI of CD14 and a high percentage of myelin-positive cells (Fig. 1d). HC macrophages, on the other hand, were defined by a high percentage of CD14^-^ and CD14^-^/CD16^-^ populations (Fig. 1d). Examining the distribution of the individual samples for these variables (Fig. 1e, Fig. S2b), it is evident that the importance of each variable varies between states, but overall a difference can be seen regardless of how the cells have been stimulated.

This difference between HC and MS samples was independent of patient sex, age, MS Severity Score (MSSS) and disease duration (Fig. S3a-d). The effect was not due to any treatment taken by the patient at the time of sampling, as untreated patient samples were equally different from HC samples compared to treated patients (Fig. S3e). Although the patient samples were collected through a cohort of sibling pairs with MS, patients from the same families did not show intra-pair similarity and were therefore considered as independent samples (Fig. S3f).

While examining each state individually provides insight into the function of the macrophages, comparing the differences between two states can give an idea of how strongly the cells respond to activating stimuli. To provide a proxy of the pro- and anti-inflammatory response, we compared cells exposed to pro- or anti-inflammatory stimuli to the homeostatic-like state. To do so, we performed a second MGCCA in which the values of the M_GM-CSF_ samples were subtracted from the M_IFNγ+LPS_ and M_IL4_ samples (Fig. 1f). Here, we again see a shift between MS and HC (Fig. 1f) but this difference is primarily present in the pro-inflammatory activation stimuli IFNγ+LPS (Fig. 1h, Fig. S2c). In this case, we see maintained levels of phagocytic cells (both myelin-positive and myelin-high) and of the CD16^+^ and CD14^+^CD16^+^ populations in MS compared to HC (Fig. 1i, Fig. S2c). This suggests a less responsive phenotype to pro-inflammatory stimuli, with a weakened anti-phagocytic response in MS macrophages. The only variable that shows MS macrophages reacting more strongly than HC macrophages is the percentage of CD14^-^CD16^+^ cells – this percentage increases in MS M_IFNγ+LPS_, a difference that is not seen in HC.

In both absolute and relative values, we thus see a higher level of phagocytosis in MS macrophages in cells treated with pro-inflammatory stimuli. The differences in surface markers are reflected in both CD14 and CD16 expression, with an over-representation of the CD14++CD16+ cells in MS patients, despite this difference not being seen in the original isolated monocytes.

### MS patient macrophages transcriptomic profile differs from HC macrophages

In order to study the complete activation profiles of the macrophages, transcriptomic profiles were analyzed through RNASeq in samples generated as above from 28 MS patients and 11 HC. Principal component analyses (PCAs) were performed both with and without consideration for the three activation stimuli, by studying each sample as one observation or by concatenating the three activation states (considering one gene in three activation states as three variables). In the concatenated PCA, which combines all samples from one individual, HC samples clustered mostly on one end of the first principal component (PC1, 22% of variance) while the MS samples showed more heterogeneity with many samples being on the opposite end of PC1 (Fig. 2a, HC: black triangles, MS: white squares). The mean value on PC1 was significantly different between MS and HC (no overlap in 95% confidence intervals calculated by boot-strapping) meaning that presence of disease was an important contributor to the transcriptomic differences between samples. This separation was further strengthened when including PC2 (14% of variance). Considering these two PCs, at least 36% of the total inter-individual variance was linked to the disease, suggesting significant large-scale transcriptomic differences between MS patients and HC.

**Fig. 2.**
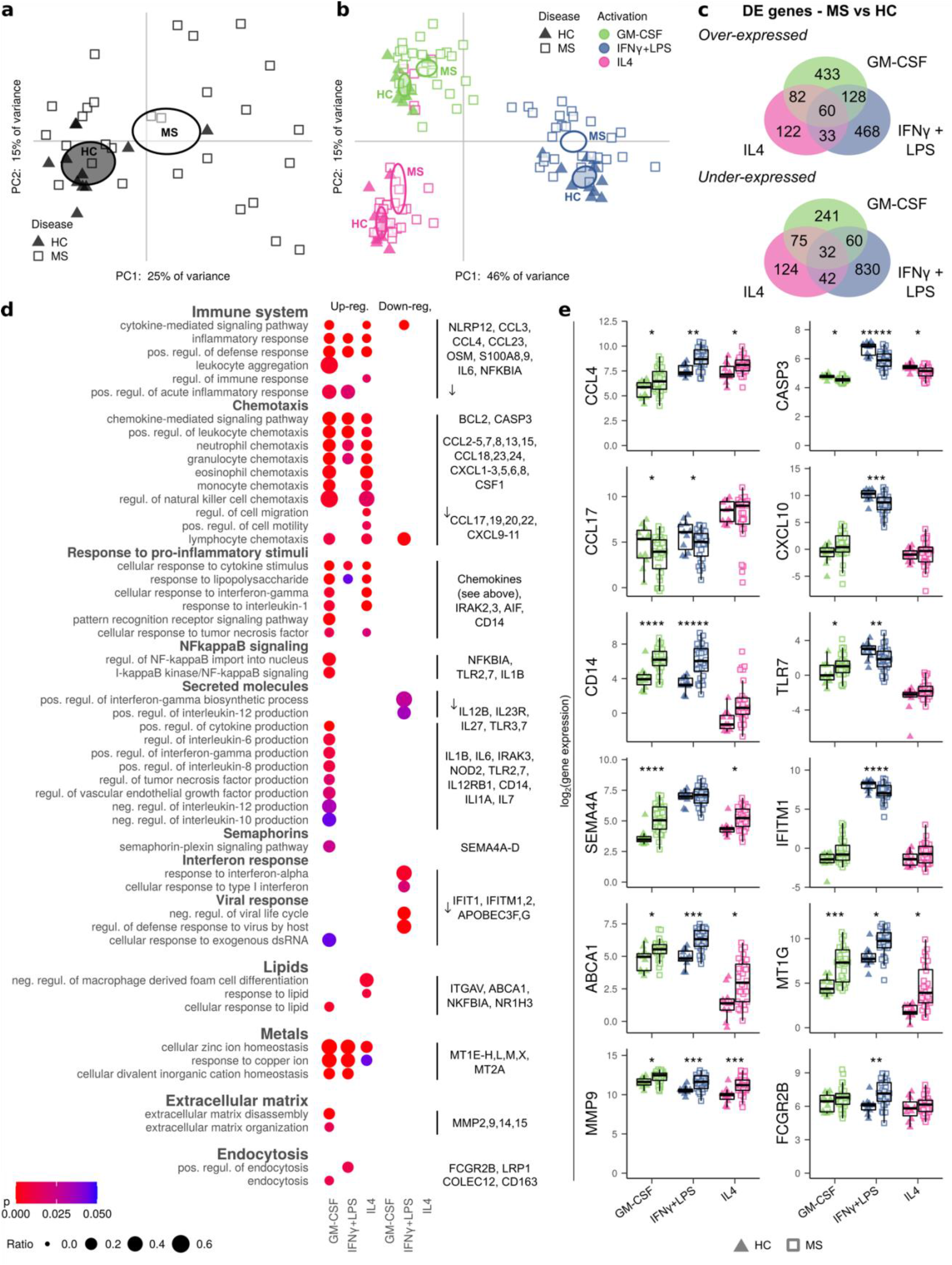
MS macrophages show a more pro-inflammatory transcriptomic profile compared to HC. **a)** PCA of all individuals, with each observation representing one individual (HC: triangles; MS: squares), and variables representing each gene in each activation state. All genes expressed in >80% of samples was included. Ellipses indicate 95% confidence interval of the group mean based on bootstrapping. **b)** PCA of the same genes as in fig. 3A, however, each observation representing one sample from one individual (HC: triangles; MS: squares; GM-CSF: green; IFNγ+LPS: blue; IL4: pink). Ellipses indicate 95% confidence interval of the group mean based on bootstrapping. **c)** Venn diagrams showing the number of differentially over- and under-expressed genes (absolute log2(fold-change)>0.5, q<0.05) between HC and MS samples in each activation state. **d)** A selection of GO terms, sorted by broader functionality, with dots indicating a significant over-representation in a given set of DE genes (adjusted p<0.05). The proportion of genes from the term and the p-value are indicated by the size and color of the dots, respectively. Examples of DE genes from at least one list of genes are provided on the right. **e)** Sample distribution for a subset of genes highlighted in d. Expression is given in log2 and grouped by activation state and disease as in b. *q<0.05, **q<0.01, ***q<0.001, ****q<0.0001, *****q<0.00001 in limma DE analysis. GO: gene ontology; HC: healthy controls; MS: MS patients (HC n=11; MS n=28).

When comparing the transcriptomic profiles of each sample (activation state), the two first principal components (58% of variance) primarily showed the effect of the activating stimuli (Fig. 2b). The three activation states were clearly separated, with the exception of five MS M_IL4_ which clustered with M_GM-CSF_. However, a difference could also be seen between MS and HC samples. There was again overlap between the groups, but the means of each group were significantly different in all activation states when considering both principal components (no overlap in 95% confidence intervals calculated by boot-strapping). Thus, the inter-sample variation was partially explained by the disease, albeit less so than by activation stimuli.

Like the functional results, the transcriptomic global differences were independent of patient sex, age, MSSS, disease duration, treatment and sibling effects (Fig. S4).

### MS patient macrophages show a globally pro-inflammatory transcriptomic profile

Differential expression (DE) analyses were performed comparing MS and HC samples in each activation state independently (log2 fold-change ≥0.5, q<0.05). In M_GM-CSF_, M_IFNγ+LPS_, and M_IL4_, respectively, 703, 689 and 297 genes were over-expressed in MS whereas 408, 964 and 273 genes were under-expressed. Several genes were differentially expressed in more than one activation state (Fig. 2c).

By analyzing the lists of DE genes for over-represented Gene Ontology (GO) terms, we identified a number of altered inflammatory pathways in all activation states (Fig. 2d). Several chemokine-coding genes, such as *CCL4* (Fig. 2e), were increased in multiple activation states, suggesting an increased capacity to attract other pro-inflammatory immune cells. A smaller number of chemokines were under-expressed in multiple states, such as *CCL17* (Fig. 2e), known to attract T helper (Th) 2 cells and typically expressed by anti-inflammatory macrophages. A number of genes important for pro-inflammatory stimuli, such as *CD14* and *TLR7* (Fig. 2e), were up-regulated in one or more states. In addition, among the most highly up-regulated genes in all states were metallothionein genes (*MT1G*, Fig. 2e). Differences were also seen in apoptosis-related genes such as *CASP3* (Fig. 2e), most notably in M_IFNγ+LPS_.

Other GO terms were primarily over-represented in one state. In M_GM-CSF_, the DE genes were consistently indicative of a pro-inflammatory profile, with alterations of genes related to cytokine secretion, semaphorins (*SEMA4A*, Fig. 2e) and extracellular matrix disassembly (*MMP9*, Fig. 2e). Different genes relating to endocytosis (*FCGR2B*, Fig. 2e) were over-expressed in MS M_GM-CSF_ and M_IFNγ+LPS_.

M_IL4_ showed a profile similar to that of M_GM-CSF_, albeit with fewer significant genes and GO terms. The only terms that were specific to M_IL4_ concerned response to lipid, with an up-regulation of genes such as *ABCA1* (Fig. 2e). However, this over-expression was not specific to M_IL4_, rather, the smaller total number of significant genes permitted the lipid-response genes to emerge as significantly over-represented.

On the other hand, M_IFNγ+LPS_ showed a state-specific down-regulation of a subset of inflammatory genes, including chemokines such as *CXCL9, 10* and *11* (*CXCL10*, Fig. 2e), known to recruit Th1 cells, certain receptors such as *TLR3* and *7* (*TLR7*, Fig. 2e), and interferon response genes of the IFIT and IFITM families (*IFITM1*, Fig. 2e).

Overall, while M_IFNγ+LPS_ showed the broadest dysregulation of genes in the number of significant DE genes (Fig. 2c), the differences in M_GM-CSF_ appeared more cohesively indicative of a pro-inflammatory state (Fig. 2d). The combination of both over- and under-expression of pro-inflammatory genes in M_IFNγ+LPS_ implies a mixed phenotype with a partially incorrect response to pro-inflammatory stimuli.

### MS patients exhibit a weaker response to pro-inflammatory stimuli compared to HC macrophages

Having seen a weakened functional response to pro-inflammatory stimuli in MS cells, and the differences in transcriptomic results between activation states, we hypothesized that MS macrophages may have a disturbed transcriptomic response to pro-inflammatory stimuli. We therefore performed inter-activation state analyses on the transcriptomic data as follows.

First, the overall similarity between two samples from the same individual was estimated using the Pearson’s correlation coefficient, calculated across all expressed genes between each pair of stimuli. The coefficients were significantly higher in MS than HC when comparing both M_GM-CSF_ with M_IFNγ+LPS_ (p<0.01, Mann-Whitney U test), and M_IFNγ+LPS_ with M_IL4_ (p<0.05, Mann-Whitney U test) (Fig. 3a), indicating a higher similarity between the activation states. This suggests a reduced response to IFNγ+LPS in MS, corroborating the differences noted above.

**Fig. 3.**
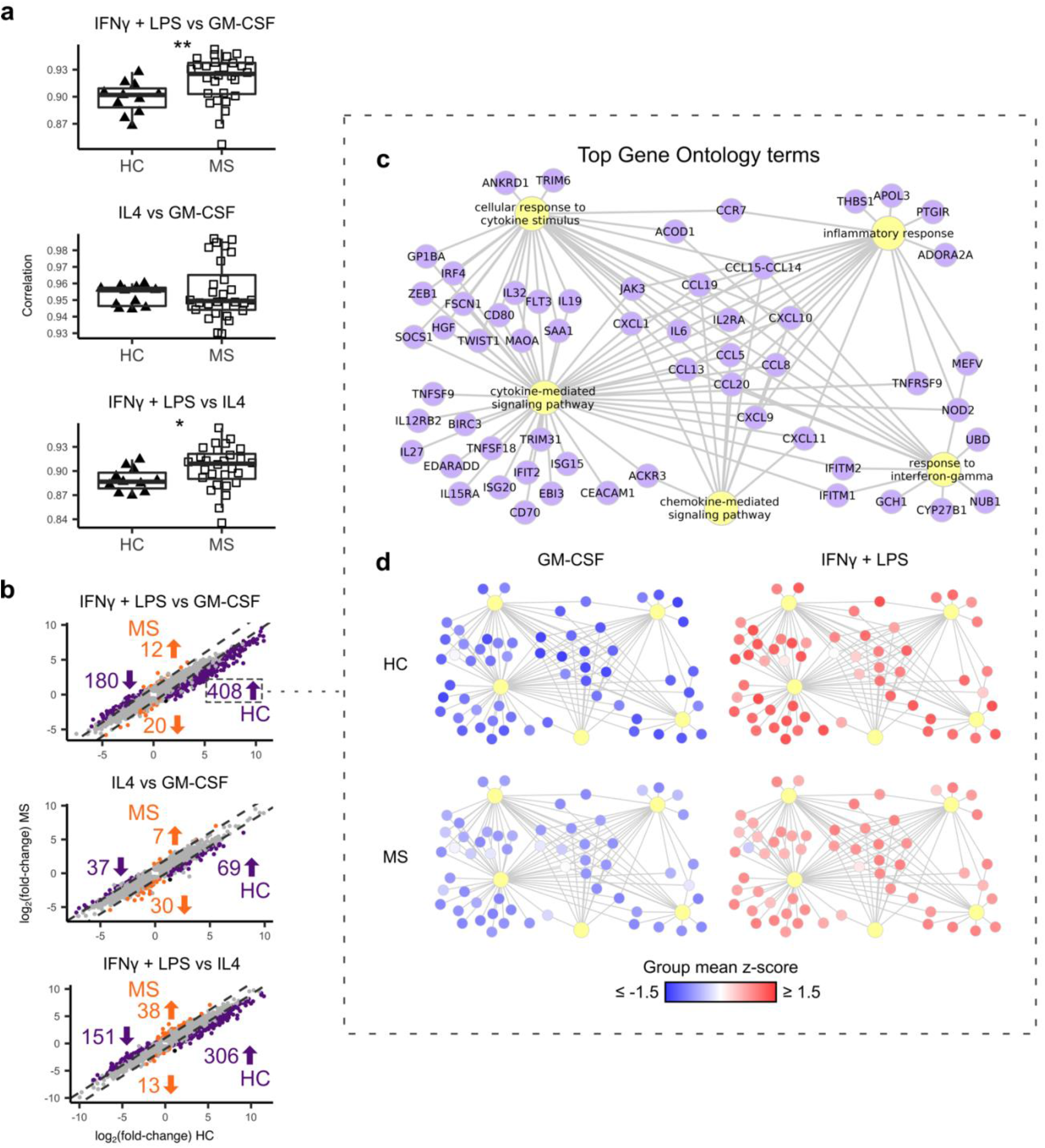
MS macrophages present a limited amplitude of response when stimulated with IFNγ+LPS. In order to compare how the transition occurs in HC and MS patients from one activation state to another we compared each pair of samples **a)** Boxplots of Pearson’s correlation coefficient between gene expression in two activation states (IFNγ+LPS vs GM-CSF, IL4 vs GM-CSF and IFNγ+LPS vs IL4), calculated for each individual (HC: triangles; MS: squares). **b)** Scatterplots showing log2(fold-change) between two states in HC (x-axis) and MS (y-axis) for each gene that is DE between the two states in MS and/or HC. Genes with a difference in mean and median log2(fold-change) greater than 0.5 between HC and MS are highlighted in orange (larger absolute fold-change in HC) and purple (larger absolute fold-change in MS). Values represent numbers of genes specific to each group and regulation (up or down). **c)** Top 5 over-represented gene ontology terms in the 408 genes with stronger up-regulation in HC than MS when comparing M_IFNγ+LPS_ to M_GM-CSF_, with the differentially regulated genes connected to their respective GO terms. **d)** Network from c colored by z-score ((mean_group_-mean_total_)/standard deviation_total_) for the mean gene expression in each activation state and disease group, with higher expression noted in red and lower expression in blue. HC: healthy controls; MS: MS patients (HC n=11; MS n=28). *p<0.05, **p<0.01 in Mann Whitney U test.

To understand whether this difference could represent an alteration of the inflammatory status of the cells, we next examined which genes caused this global difference. DE analyses were performed for each pair of stimuli, in MS and HC samples separately (6 analyses in total). For all the DE genes between two states (significantly different in HC and/or MS), we plotted the fold-change in MS samples against the fold-change in HC samples (Fig. 3b). In all comparisons, a larger number of genes were more strongly regulated in HC (purple dots), than MS (orange dots) (difference in mean and median log_2_ (fold-change) > 0.5), again suggesting a weaker response in MS. Like the difference in correlation, this effect was most noticeable in the difference between M_IFNγ+LPS_ and M_GM-CSF_, with 408 and 180 genes being more strongly up- and down-regulated, respectively, in HC compared to MS. In contrast, only 32 genes were more strongly regulated in MS. In the 408 genes that were more strongly up-regulated in the HC, the five most over-represented GO terms were inflammatory response, cellular response to cytokine stimulus, cytokine-mediated signaling pathway, chemokine-mediated signaling pathway and response to interferon-gamma (Fig. 3c), again implying a partially reduced inflammatory response to IFNγ+LPS in MS. These genes include the above identified chemokines *CXCL9-11* and interferon response genes of the IFIT and IFITM families (Fig. 3c).

To ensure that this reduced response was not a result of saturation of pro-inflammatory genes already over-expressed in M_GM-CSF_, the mean expression level of these genes were visualized for each group of samples (M_GM-CSF_ and M_IFNγ+LPS_ in HC and MS) as a z-score (group mean subtracted by total mean, divided by the standard deviation for each gene) in Fig. 3d. As should be the case, we can see that all genes were more strongly expressed in HC M_IFNγ+LPS_ than M_GM-CSF_ (Fig. 3d, upper panels). However, despite the expression levels of most genes already being slightly higher in MS M_GM-CSF_ compared to HC, the expression was lower in MS M_IFNγ+LPS_ than in HC (Fig. 3d, lower panels), arguing against a “saturation” effect and for a deviant response and altered final state. Thus, our data speak to a dysregulated response to pro-inflammatory stimuli beyond the altered state seen in homeostatic-like cells.

### Co-expressed genes involved in migration, apoptosis and metabolism are dysregulated in MS macrophages

By investigating transcriptomic differences in a modular way, rather than gene by gene, it is possible to identify groups of co-expressed genes that each show a small intergroup difference but together likely have a larger effect. To this end, we performed a weighted gene co-expression network analysis (WGCNA). In order to avoid the differences between activation states masking the effect of the disease, we constructed the network by calculating the correlation between each pair of genes in each activation state separately, then using the maximum correlation for each gene pair in the module construction (Fig. 4a, Fig. S5a). For each module produced in the analysis, the genes in the modules were analyzed to identify over-represented GO terms and KEGG pathways (Fig. S5b). The eigengene (first principal component) value was then calculated for each module and sample using the PCA for each activation state individually. The correlation between these eigengene values and the disease status (HC or MS) was then calculated (Fig. S5b-c). All modules with a significant correlation (FDR-adjusted asymptotic p<0.05) in at least one activation state and at least one significantly over-represented GO term or KEGG pathway are represented in Fig. 4b. The module with the strongest correlation to disease contained several chemokine and metal homeostasis genes, reiterating the differences observed in the DE analysis.

**Fig. 4.**
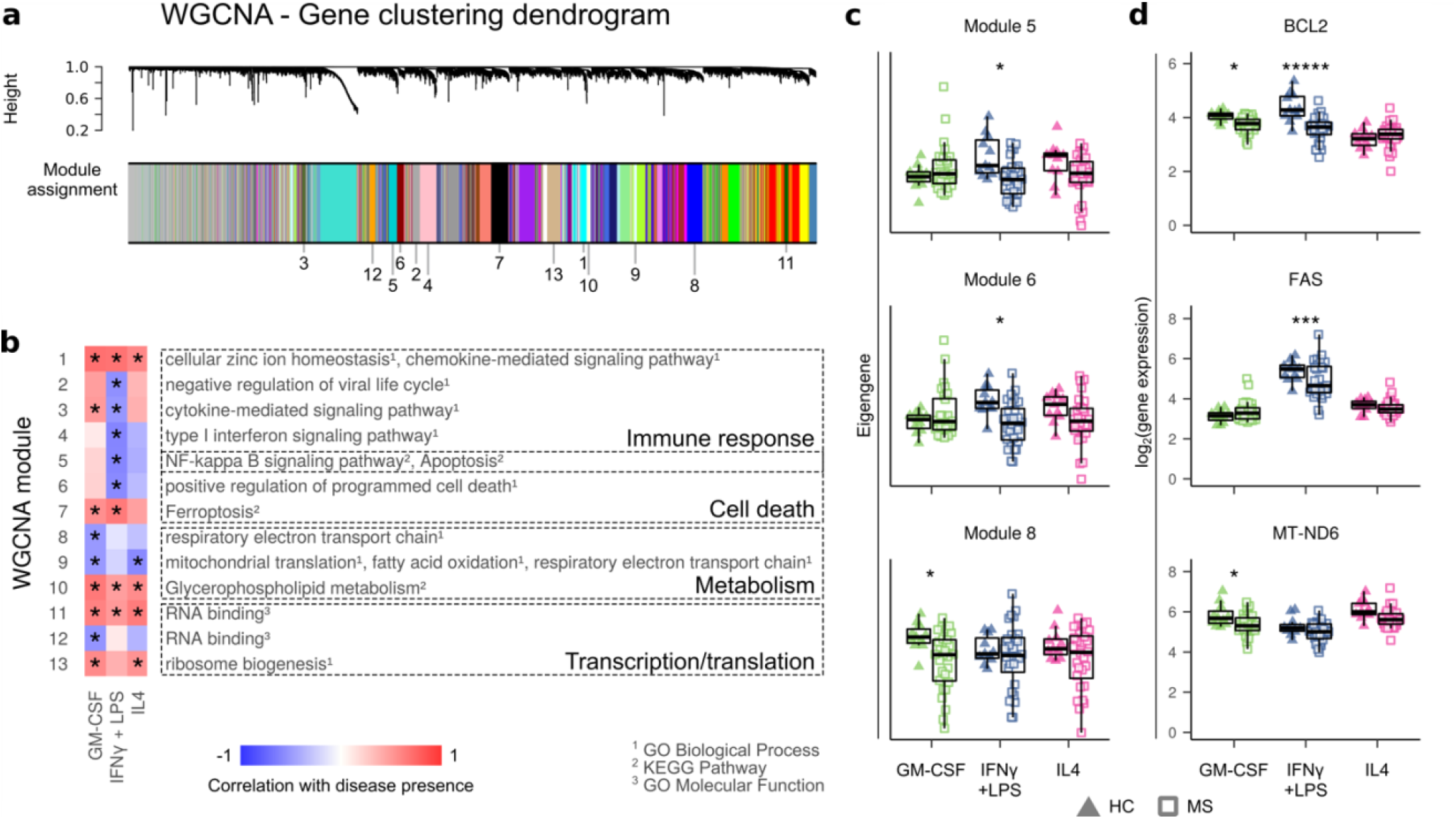
Modules of co-expressed genes involved in inflammatory and metabolic pathways show altered expression in MS macrophages compared to HC macrophages. **a)** Clustering of genes in multi-dataset WGCNA, with module assignment indicated below. Multi-dataset network was produced by comparing correlations between genes for each state individually and basing clustering on the maximum correlation for each gene pair. **b)** Heat map of correlation between presence of disease and eigengenes of each module and activation. Only modules with significant correlation and functional annotation terms are shown. Functional annotations were given by Enrichr from databases GO biological process, GO molecular function, and KEGG pathways (adjusted p<0.05). **c)** Eigengene values in three modules, for each sample and module, grouped according to disease (HC: triangles; MS: squares) and activation state (GM-CSF: green; IFNγ+LPS: blue; IL4: pink). **d)** Expression of examples of genes from each module shown in c (grouped and visualized as in c). HC: healthy controls; MS: MS patients (HC n=11; MS n=28). *p< 0.05. Correlations were calculated with FDR-corrected default WGCNA functions (b, c), differential expression was tested with limma (d).

Six modules were negatively correlated with disease in M_IFNγ+LPS_, containing many of the interferon-response genes seen previously, but also several genes involved in cell death (modules 5 and 6, Fig. 4C). However, these modules included genes that promote survival as well as cell death, such as *BCL2* and *FAS* (Fig. 4d). When comparing the levels of the genes present in the Apoptosis KEGG pathway, MS M_IFNγ+LPS_ showed both over- and under-expression of several pro- and anti-apoptotic genes (Fig. S6), complicating the interpretation of the impact on apoptosis. However, the under-expression of several caspase genes suggests a dysfunction of this important effector aspect of the pathway (Fig. S6).

Two modules with several genes involved in the respiratory electron transport chain were negatively correlated with disease in M_GM-CSF_ (modules 8 and 9, Fig. 4c and Fig. S5c). Although each gene on its own showed a relatively small dysregulation (MT-ND6, Fig. 4d), the concerted decreased expression of several genes involved in the process suggest a global reduction of oxidative metabolism, a key feature of anti-inflammatory cells [32]. For this reason, we chose to further study the metabolic phenotypes of the cells, as described below.

There were also three modules that correlated with disease and contained several genes relating to RNA binding and ribosome biogenesis (Fig. 4b), suggesting an even more complex dysregulation of mRNA and protein levels.

### MS homeostatic-like macrophages present dysregulated mitochondrial energy metabolism

To define the metabolic phenotype of the cells, a metabolomic analysis was performed using liquid chromatography mass spectrometry (LC-MS) on samples from cells prepared in the same way as for the transcriptomic analysis. As the transcriptomic changes related to metabolism were mostly noted in M_GM-CSF_, we chose to focus on this activation state. We included only untreated patients to eliminate potential effects of metabolism-modifying treatment (HC n = 6, MS n = 7), effects notably observed in DMF treated individuals [9]. In order to identify changes in energy metabolism, we examined the metabolites and genes (using the previous transcriptomic results) involved in 4 main pathways: fatty acid oxidation; the tricarboxylic acid (TCA) cycle; oxidative phosphorylation; and glycolysis (Fig. 5).

**Fig. 5.**
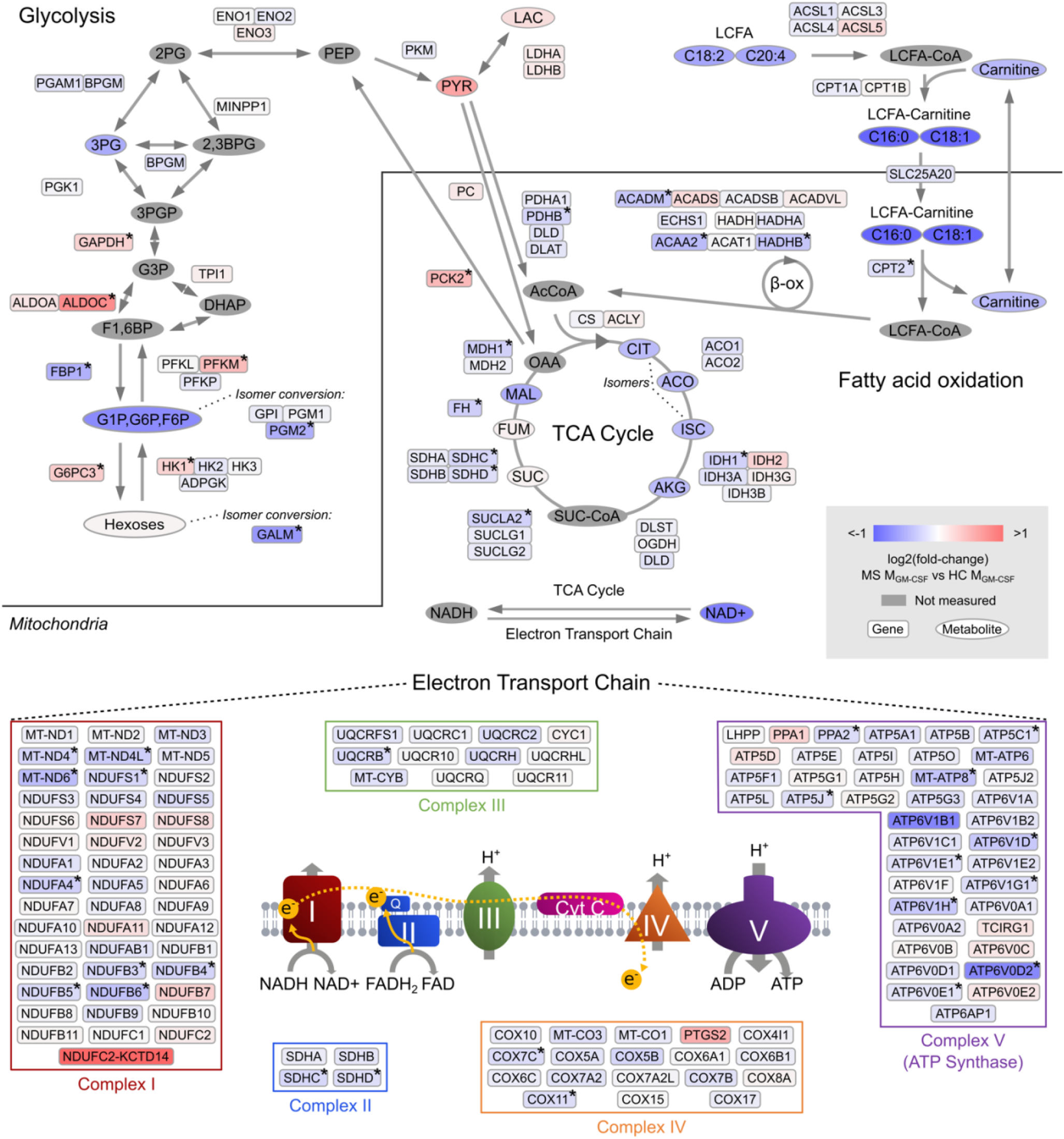
MS homeostatic-like macrophages show several signs of alteration in the TCA cycle and respiratory chain. Genes and metabolites included in the KEGG pathways Glycolysis, Fatty acid degradation, TCA cycle and Oxidative phosphorylation are organized according to pathways. Each element is colored to indicate mean abundance in MS M_GM-CSF_ relative to HC (Δlog2(mean), over-expression in red, under-expression in blue, HC n=6; MS n=7). Genes involved in oxidative phosphorylation are organized into the complexes in which their gene products take part. Genes that were not expressed in at least one sample were excluded in the figure. *significantly altered genes (q<0.05) in limma differential expression analysis. HC: healthy controls; MS: MS patients; 2,3BPG: 2,3-bisphosphoglyceric acid; 2PG: 2-phosphoglyceric acid; 3PG: 3-phosphoglyceric acid; 3PGP: 3-phospho-D-glyceroyl phosphate; AcCoA: acetyl coenzyme A; ACO: aconitate; AKG: alpha-ketoglutarate; β-ox: beta-oxidation; CIT: citrate; DHAP: dihydroxyacetone phosphate; F1,6BP; fructose-1,6-bisphosphate; FUM: fumarate; G1P: glucose-1-phosphate; G3P: 3-phosphoglycerate; G6P: glucose-6-phosphate; ISC: isocitrate; LAC: lactate; LCFA: long-chain fatty acid; MAL: malate; OAA: oxaloacetate; OXSUC: oxalosuccinate; PEP: phosphoenolpyruvic acid; PYR: pyruvate; SUC: succinate; SUC-CoA: succinyl coenzyme A.

While none of the individual metabolites included in Fig. 5 showed significant differences between patients and HC (Fig. S7), the concordance between mean differences in metabolite abundance and significant differences in gene expression nonetheless gives an idea of the activity of the pathways.

Overall, patients with MS displayed a global down-regulation of all mitochondrial pathways. Mitochondrial β-oxidation of fatty acids was down-regulated in MS samples with both (i) decreased levels of long-chain fatty acids (arachidonic and linolenic acids), acylcarnitines (oleyl- and palmitoyl-carnitines) and carnitine, as well as (ii) a significant under-expression of four major genes (*CPT2, HADHB, ACADM* and *ACAA2*) encoding carnitine palmitoyltransferase type 2, the trifunctional protein, medium-chain acyl-coenzyme A dehydrogenase and acetyl-CoA acyltransferase 2, respectively.

We also observed decreased levels (log2 fold-change <-0.4) of TCA intermediates (citrate, isocitrate, aconitate, α-ketoglutarate and malate) in MS samples, together with significant under-expression of almost all genes involved in the TCA cycle (*IDH1, SUCLA2, SDHC, SDHD, FH* and *MDH1*).

Downstream, all five complexes of the electron transport chain showed the following signs of under-expression: 1) at least one significantly under-expressed gene, 2) no significantly over-expressed genes, 3) a larger number of genes with downregulated expression (significant or not) (Fig. 5). Consistent with this observation, NAD^+^, which results from the oxidation of NADH by the electron transport chain, was less abundant in MS samples.

Conversely, levels of lactate and pyruvate tended to be higher in MS samples (Fig. 5, Fig. S7), suggesting a shift from oxidative metabolism to glycolysis. Likewise, four key genes (*HX1, PFKM, ALDOC* and *GAPDH*) involved in glycolysis and two regulators of gluconeogenesis (*PCK2* and *G6PC3*) were significantly over-expressed in patients with MS.

Collectively, despite the intragroup heterogeneity, these combined data strongly suggest a reduced capacity of oxidative metabolism in MS M_GM-CSF_, consistent with a pro-inflammatory phenotype.

## Discussion

Using peripheral monocytes from MS patients and HC, we provide evidence of a pro-inflammatory profile in MS monocyte-derived macrophages prior to lesion exposure, regardless of activating stimuli, reminiscent of trained immunity. A metabolic switch accompanies this over-inflammatory response with a downregulation of several pathways of mitochondrial energy metabolism. However, MS macrophages also show signs of tolerance toward pro-inflammatory stimulation.

### Pro-inflammatory tendencies of MS patient macrophages

MS macrophages over-expressed several cytokine genes, indicating a role in maintaining a pro-inflammatory environment and promoting immune cell infiltration into the CNS. For instance, classical pro-inflammatory interleukins *IL1B* and *IL6* were increased in MS patient macrophages. Furthermore, genes encoding for ligands of CCR2 and CCR5 receptors (CCLs 2-5, 7, 8, and 13) were significantly over-expressed in at least one activation state, suggesting an implication of macrophages in the increased percentage of CD4^+^CCR2^+^CCR5^+^ cells observed in the CSF in MS during relapse [61]. These CD4^+^CCR2^+^CCR5^+^ cells in turn produced high levels of pro-inflammatory cytokines and were reactive to myelin basic protein (MBP) [61]. In addition, CCL2 and CCL5, which were both over-expressed in MS conditions, induce stronger *in vitro* migratory capacities in monocytes from MS patients than from HC [18], implicating macrophages in increased recruitment of infiltrating monocytes. Interestingly, MS patient macrophage under-expressed *CCL17* which is known to attract Th2 and regulatory T cells, subtypes of anti-inflammatory lymphocytes [73]. In total, these transcriptomic data suggest that infiltration of disease-associated cell types could be exaggerated by macrophage defects observed in MS conditions.

Another family of genes that was over-expressed in MS macrophages of all activation states was MTs, which are cysteine-rich proteins capable of binding metals. They are important for copper and zinc homeostasis, protection against oxidative stress and sequestration of heavy metals [45, 60]. Several roles for MT in immune regulation have been proposed [64], and MT over-expression has been described in MS CNS [52]. Beyond immune regulation, MT expression could be related to the reduced concentration of zinc observed in MS [12] through sequestration. As zinc binds to myelin proteins such as MBP and seems important for myelin structure and/or function [15, 67], its reduction may be of direct importance in the degeneration of myelin.

The identified transcriptomic signature matches a differential expression of cell surface markers. Expression of CD14 and CD16 showed significant differences between MS patient and HC with a larger proportion of CD14^++^CD16^+^cells. Interestingly, this result reflects the observation that CD16^+^ monocytes are present in MS active lesions, participating to blood brain barrier breakdown and T cells invasion of the CNS [71].

Overall, we thus see that macrophages derived from peripheral monocytes of MS patients present a phenotype that is both reflective of known characteristics in MS lesions, and that indicate a role of macrophages in important pathological events. As such, this model shows potential to aid identification of biomarkers as well as testing of treatments targeting the innate immune system, a strategy so far under-utilized in MS treatment.

### MS patient macrophages exhibit a perturbed energy metabolism

The activation as pro- and anti-inflammatory macrophages relies on specific metabolic profile, with increased anaerobic glycolysis and oxidative metabolism, respectively [32]. We observe that MS patient macrophages, in absence of pro-inflammatory signals (M_GM-CSF_), exhibit a preferential glycolytic metabolism when compared to HC macrophages, thus already showing signs toward a pathogenic phenotype. Interestingly, perivascular macrophages with high glycolytic capacity were identified in an MS animal model and showed increased transmigratory functions [30], implicating this metabolic dysregulation in immune cell infiltration in MS. The same study showed indications of similar cells in post-mortem MS tissue.

Lactate, another indicator of aerobic glycolysis and mitochondrial dysfunction that was more prevalent in MS patient macrophages, is also detected at higher levels MS cerebrospinal fluid [2] and serum [3].

Here, enhanced glycolysis is accompanied by a reduction of electron transport chain gene expression in MS patient macrophages, an observation that was also reported on a protein level in MS patient lymphocytes [23]. These mitochondrial alterations observed in immune cells would have therefore an impaired redox status and anti-oxidant capacities. In particular, reactive oxygen species (ROS) are believed to be a main instigator of oligodendrocyte alterations and neurotoxicity [1]. ROS production is linked to energy metabolism [55], and the reduced mitochondrial metabolism seen here in MS macrophages could thus reflect a destructive phenotype contributing to neural damage.

In addition to the changes in the electron transport chain, we saw disturbances in the TCA cycle and fatty acid metabolism, indicating overall reduction of oxidative metabolism in MS. Oxidative metabolism is both a hallmark of and necessary for the anti-inflammatory state in macrophages [25, 68]. The observed imbalance between glycolysis and oxidative metabolism may thus be a result and/or a cause of the pro-inflammatory MS phenotype even in the absence of pro-inflammatory signals.

A few metabolic treatment strategies have been proposed to reduce degeneration in neural cells in progressive MS [24]. It is however recognized that many metabolic therapeutics could act on macrophages as well [53] and more specifically that a metabolic switch (from aerobic glycolysis to oxidative phosphorylation) is essential for promotion of a pro-regenerative states. Interestingly, one of the most common first line treatments in MS, dimethyl fumarate, is believed to function through metabolic alterations [31, 53], although it is unclear which cell types are primarily implicated in this correction.

### Trained innate immunity

The concept of trained innate immunity describes the increased reactivity to novel signals after a prior activation. Increased pro-inflammatory cytokine production and altered metabolism are both hallmarks of this second reactive state [49]. Although monocytes are short-lived, the trained immunity is epigenetically programmed in the bone marrow and can thus reflect previous exposure to antigens from a time prior to the generation of the tested monocytes [29]. Our work shows that pro-inflammatory tendencies of MS patient macrophages exist well before they enter the highly inflammatory environment of a lesion just after exposure to GM-CSF.

Trained immunity has previously been identified as a phenomenon through which the innate immunity could contribute to inflammatory diseases [5], participating both to disease initiation and maintenance or aggravation of deleterious environment. For example, in atherosclerosis and systemic lupus erythematous, monocytes/macrophages share several features observed in MS patient macrophages such as increased expression of pro-inflammatory cytokines [8, 48], metabolic rewiring [8, 27], and overrepresentation of pro-inflammatory CD14^+^CD16^+^ cells [48, 58].

However, the manifestation of innate immune memory can also be disease specific. Here we document yet another MS macrophage peculiarity. In MS patient macrophages, we did not observe the expected further increased response when MS macrophages were exposed to IFNγ+LPS, to the point where a subset of genes involved in inflammatory responses showed lower expression in MS M_IFNγ+LPS_ than HC M_IFNγ+LPS_. This is reminiscent of previously described LPS tolerance in macrophages, in which interferon response gene expression is reduced upon a second stimulation [43]. LPS exposure is an unlikely cause of MS patient-specific phenotypes, but a similar desensitization might be triggered by other causes. While this desensitization intuitively would seem beneficial to the patient, it is also possible that this incomplete activation fails to activate important inflammatory resolution-inducing pathways. For instance, death of pro-inflammatory cells is a regulatory phenomenon described generally in resolution [62] and specifically in myelin regeneration [42]; the failure to up-regulate apoptosis effector genes in MS M_IFNγ+LPS_ could result in a longer-lived pro-inflammatory macrophage population and failure to institute a pro-regenerative one.

However, it is important to note that while the response overall was weaker in MS macrophages, MS M_IFNγ+LPS_ still showed up-regulation of several pro-inflammatory cytokines compared to HC, suggesting a mixed phenotype with features of tolerance and trained immunity. We also see a mixed pro- and anti-inflammatory phenotype in MS MIL-4, with anti-inflammatory genes weakly up-regulated. These phenotypes may be related to the intermediate activation state that has previously been described in MS lesions [69].

In conclusion, our data imply a perturbed macrophage response with characteristics of both trained innate immunity and tolerance. Therefore, we propose that the predisposition to a pro-inflammatory state, as well as the response to activating stimuli, must be considered in an MS-specific context when predicting how the innate immune system can be targeted in treatment.

### Concluding remarks

This study implicates a role of monocyte-derived macrophage function and transcriptome as mediators and/or biomarkers of MS. In an artificial environment, we were able to mimic several aspects of MS lesions, supporting the role of innate immune system memory in the inflammatory lesion environment. For a better understanding of how this dysregulation affects different patient groups, future work would likely benefit from connecting cellular biology and clinical observations through more extensive phenotyping, in particular keeping in mind the potential functions of the cell in disease.

## Acknowledgments

We are grateful to all the patients and the volunteers that participated to this study, together with the members of the Zujovic/Nait Oumesmar team for their support. We are indebted to the Bouvet-Labruyère family and the OCIRP foundations for their constant support to MS research at the ICM. The authors acknowledge Dr Michel Mallat and Dr. Jaime de Juan-Sanz for their critical reviews of the work. The authors wish to thank the cell culture (CELIS), sequencing (i-GENSEQ), biostatistics facilities (ICONICS) but also the fundraising, scientific affairs and the administrative department of the Paris Brain Institute. We want to thank also Sarah Taieb Tamacha from the Paris Brain Institute-CIC for her efficient management of MS patient recruitment of the MS-BIOPROGRESS cohort and for supervision of blood sampling from MS patients.

## Funding

This work was supported by the OCIRP foundation, Bouvet Labruyère Price, SANOFI Innovation Awards program (iAwards) 2018 and the program “Investissements d’Avenir” ANR-10-IAIHU-06,“Translational Research Infrastructure for Biotherapies in Neurosciences” ANR-11-INBS-0011–NeurATRIS and “Idex Sorbonne Université dans le cadre du soutien de l’Etat aux programmes Investissements d’Avenir”.

## Author contributions

VZ designed and supervised the project. JF performed data acquisition and analysis, and drafted and wrote the manuscript. VZ wrote, reviewed and edited the manuscript. CL supervised the MSBIOPROGRESS cohort management and part of the project. BF supervised part of the project. CL, BS, EM participated in patient recruitment. CB, LGN, FD, AG, FI, MP participated to data acquisition and analysis. FM supervised part of the project and data analysis. AT supervised statistical analysis. All authors discussed the results and commented on the manuscript.

## Competing interests

Dr Stankoff reports grants and personal fees for lectures from ROCHE, SANOFI-GENZYME, and MERCK-SERONO, personal fees for lectures from NOVARTIS, BIOGEN and TEVA, all outside the submitted work.

Dr Maillart reports grants and personal fees from Biogen, Novartis, and Roche; and personal fees from Merck-Serono, Teva, Sanofi-Genzyme, and Ad Scientiam outside of the submitted work.

Dr Louapre has received consulting or travel fees from Biogen, Novartis, Roche, Sanofi, Teva and Merck Serono, and research grant from Biogen, none related to the present work.

Dr Zujovic received SANOFI Innovation Awards program (iAwards) 2018, related to this work

### Abbreviations

CNS: central nervous system
DE: differential expression / differentially expressed
EDSS: expanded disease severity score
FBS: fetal bovine serum
GM-CSF: granulocyte macrophage colony-stimulating factor
GO: gene ontology
HC: healthy control
IFN: interferon
IL: interleukin
LPS: lipopolysaccharide
MGCCA: multiway regularized canonical correlation analysis
MFI: mean fluorescence intensity
MS: multiple sclerosis
MSSS: multiple sclerosis severity score
MT: metallothionein
OPC: oligodendrocyte precursor cell
PBMC: peripheral blood mononuclear cell
PCA: principal component analysis
ROS: reactive oxygen species
TCA: tricarboxylic acid
Th: T helper
WGCNA: weighted gene co-expression network analysis

