## Supplementary Figures for "Dysregulated functional and metabolic response in multiple sclerosis patient macrophages correlate with a more inflammatory state, reminiscent of trained immunity"

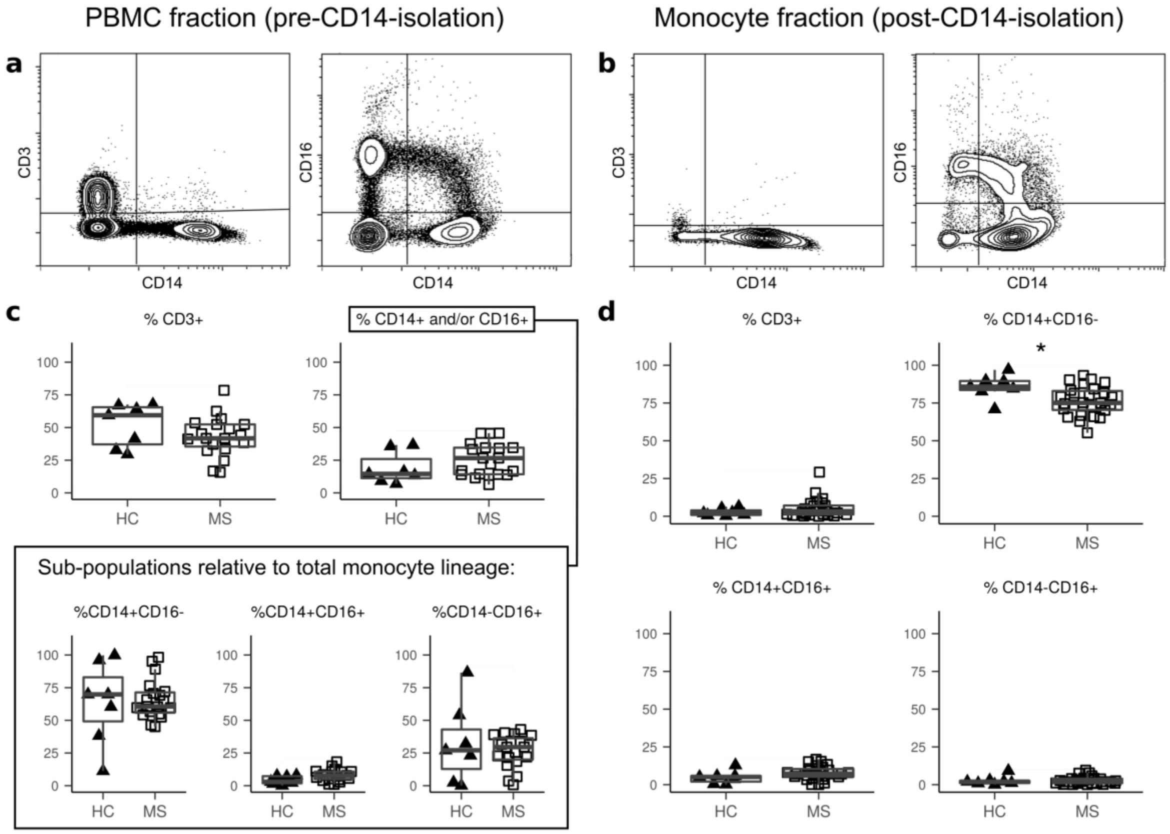


**Fig. S1** PBMCs show no significant differences in CD3, CD14 and CD16 expression between MS patients and HC. a) Representative dotplots of CD3, CD14 and CD16 expression in PBMCs prior to $CD14^+$ selection. b) Representative dotplots of CD3, CD14 and CD16 expression in $CD14^+$ population. c) Percentages of $CD3^+$ and $CD14^+/CD16^+$ populations in PBMCs prior to $CD14^+$ selection. Values in the second row of graphs have been calculated as a proportion of the total number of cells positive for CD14 and/or CD16 (HC n=7; MS n=19). d) Percentages of $CD3^+$ and $CD14^+/CD16^+$ populations in monocytes after $CD14^+$ selection (HC n=7; MS n=26). HC: healthy controls; MS: MS patients. *p<0.05, Mann Whitney U test.

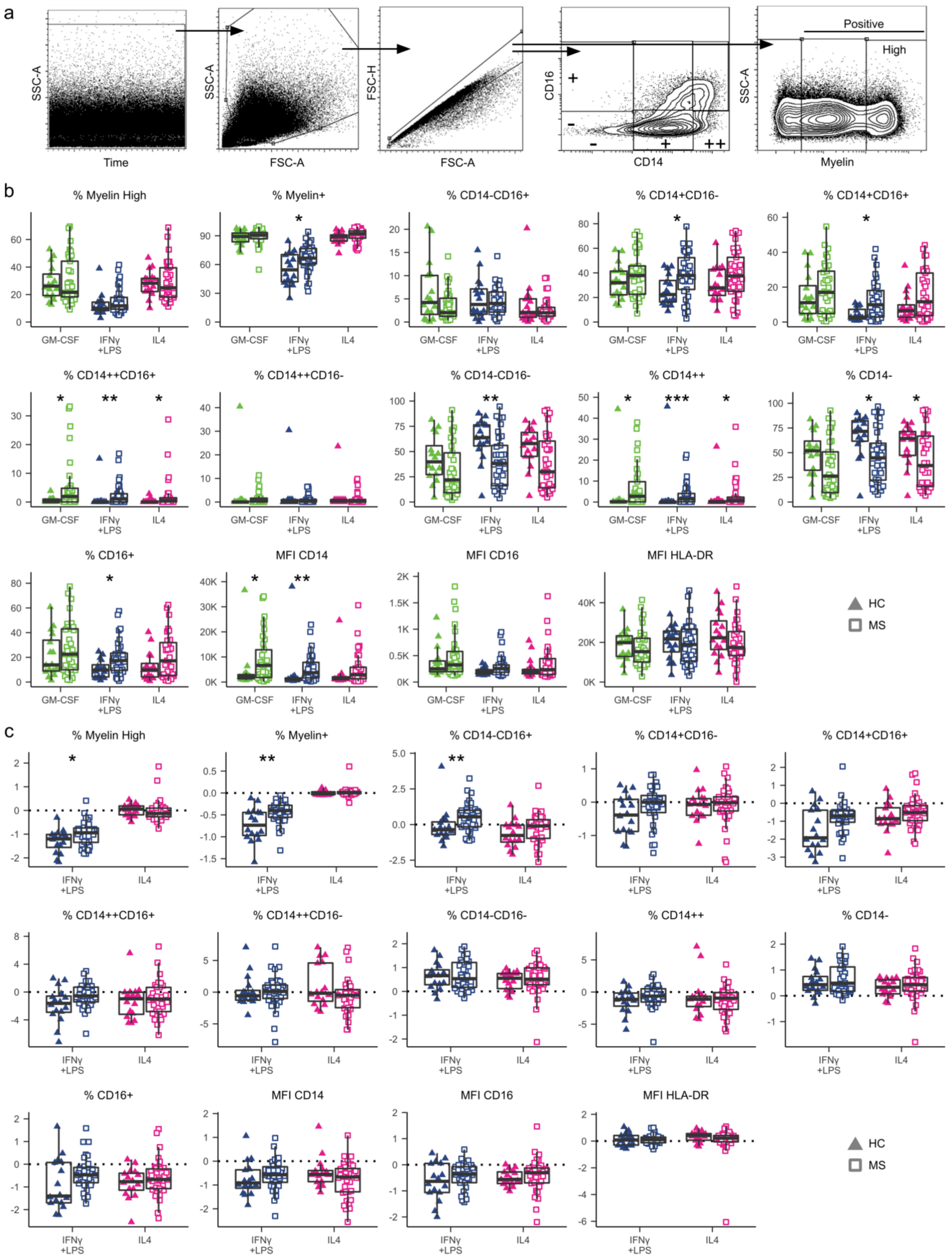
a
SSC-A
Time
SSC-A
FSC-A
FSC-H
FSC-A
CD16
+
-
-
+
++
CD14
SSC-A
Positive
High
Myelin
b
% Myelin High
% Myelin+
% CD14-CD16+
% CD14+CD16-
% CD14+CD16+
% CD14++CD16+
% CD14++CD16-
% CD14-CD16-
% CD14++
% CD14-
% CD16+
MFI CD14
MFI CD16
MFI HLA-DR
GM-CSF
IFNγ +LPS
IL4
HC
MS
c
% Myelin High
% Myelin+
% CD14-CD16+
% CD14+CD16-
% CD14+CD16+
% CD14++CD16+
% CD14++CD16-
% CD14-CD16-
% CD14++
% CD14-
% CD16+
MFI CD14
MFI CD16
MFI HLA-DR
IFNγ +LPS
IL4
HC
MS

**Fig. S2** Phagocytosis of myelin and surface expression of CD14, CD16 and HLA-DR in macrophage samples (HC n=16; MS n=33). a) Gating strategy for flow cytometry analysis presented in Fig. 2. b) Distribution of samples for all variables studied in Fig. 2a-e, grouped by disease status and activation state (HC: triangles; MS: squares; GM-CSF: green; IFNγ+LPS: blue; IL4: pink). c) Same as A for all variables studied in Fig. 2f-j. HC: healthy controls; MS: MS patients. * $p < 0.05$, Mann-Whitney U test not corrected for multiple testing.

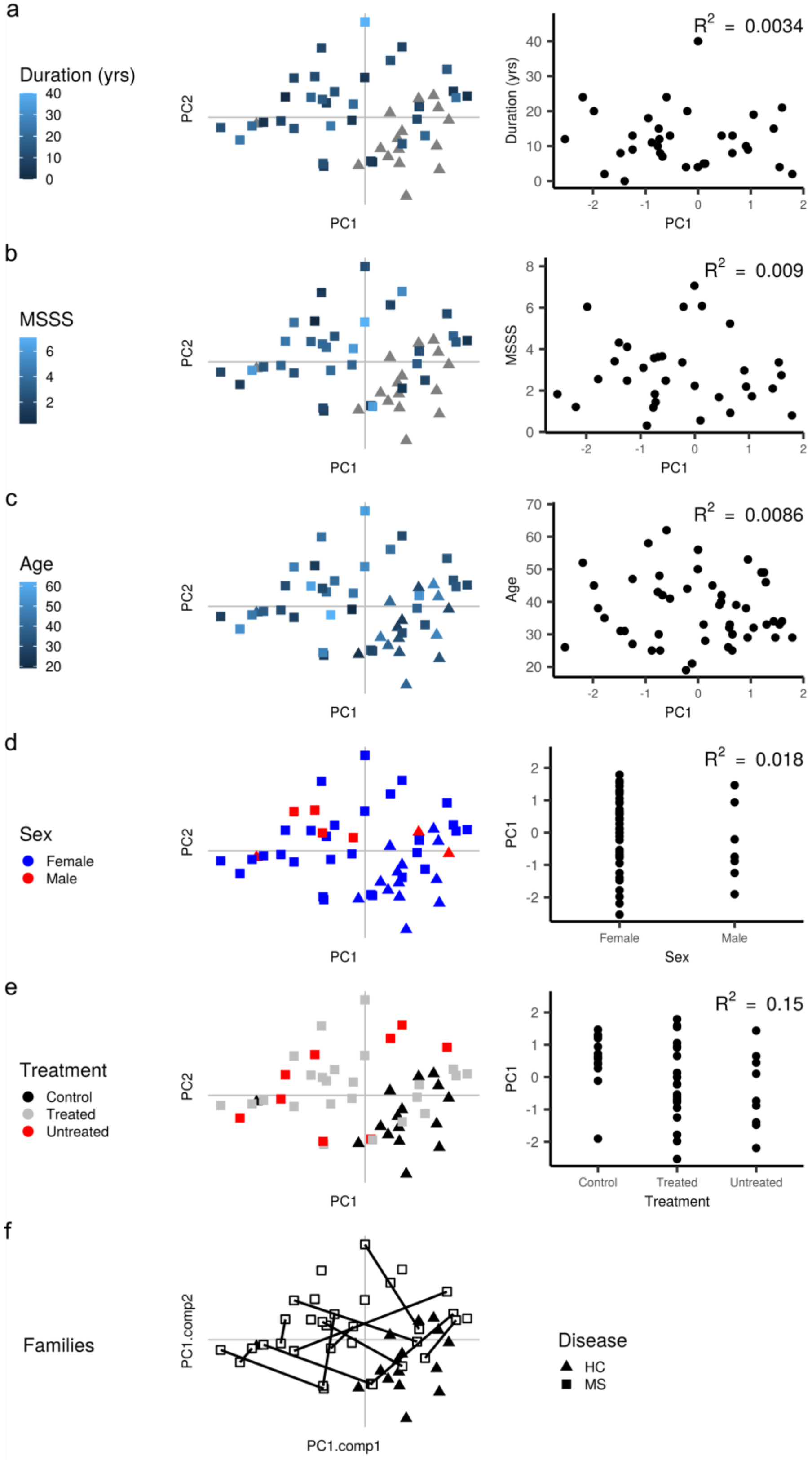
a
Duration (yrs)
40
30
20
10
0
PC2
PC1
$R^2$ = 0.0034
Duration (yrs)
b
MSSS
6
4
2
$R^2$ = 0.009
c
Age
60
50
40
30
20
$R^2$ = 0.0086
d
Sex
Female
Male
$R^2$ = 0.018
e
Treatment
Control
Treated
Untreated
$R^2$ = 0.15
f
Families
PC1.comp2
PC1.comp1
Disease
HC
MS

**Fig. S3** Flow cytometric data do not correlate with potential confounders. a) Graph in Fig. 2B with data points colored according to patient disease duration (longer duration: light blue; shorter duration: dark blue) (left), and plot showing correlation between disease duration and value on PC1 (right), as this component represents the largest difference between HC and MS. b-e) Same as (a) with MSSS (more severe score: light blue; less severe score: dark blue), age (older: light blue; younger: dark blue), sex (female:blue; male: red) and treatment status (HC: black; treated MS: grey; untreated MS: red) as potential confounding factors. f) Graph in Fig. 2b with lines drawn between data points representing pairs of siblings. HC: healthy controls; MS: MS patients; MSSS: MS Severity Score.

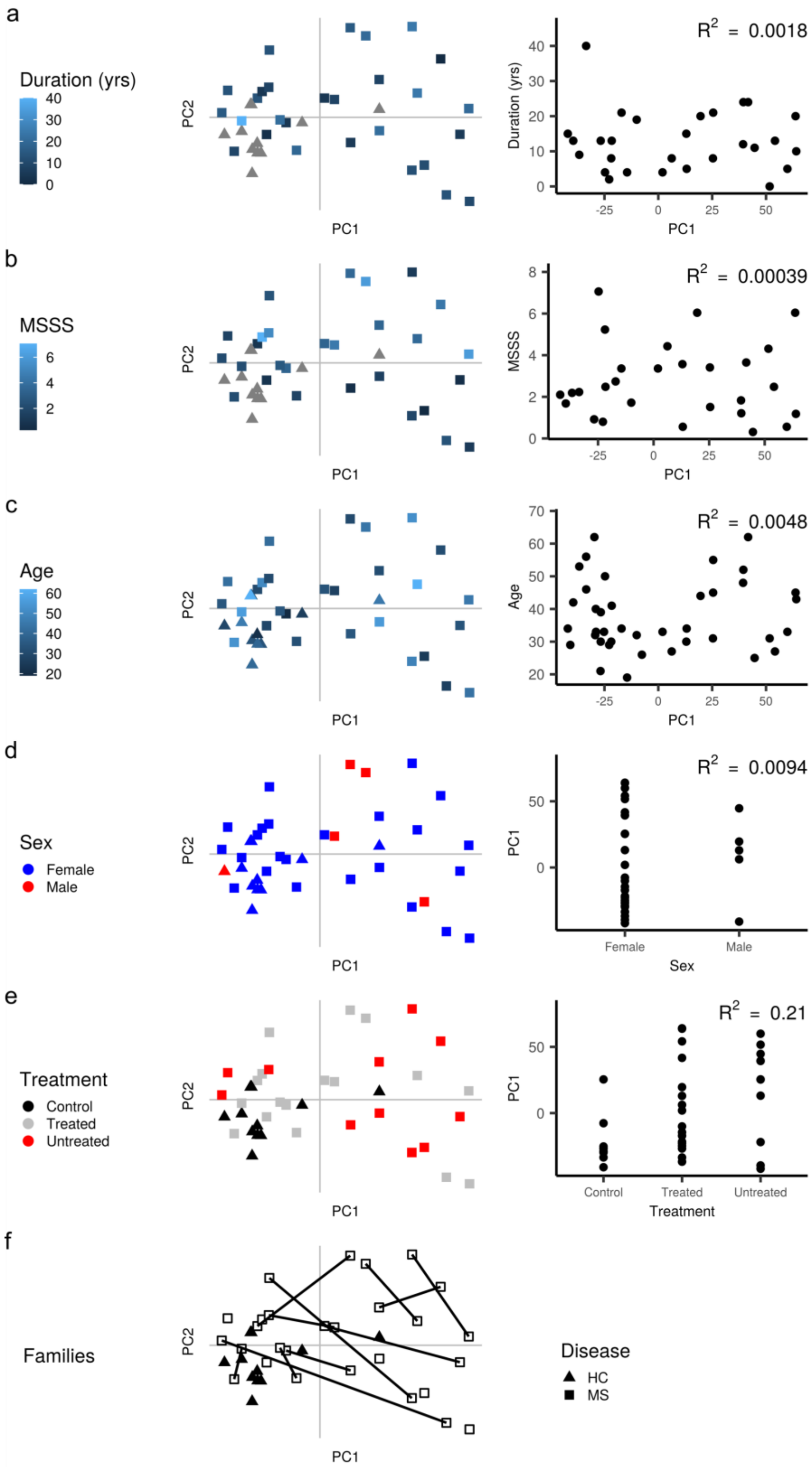
a
Duration (yrs)
40
30
20
10
0
PC2
PC1
$R^2$ = 0.0018
Duration (yrs)
-25
0
25
50
b
MSSS
6
4
2
$R^2$ = 0.00039
8
c
Age
60
50
40
30
20
$R^2$ = 0.0048
70
d
Sex
Female
Male
$R^2$ = 0.0094
Female
Male
e
Treatment
Control
Treated
Untreated
$R^2$ = 0.21
f
Families
Disease
HC
MS

**Fig. S4** Transcriptomic data do not correlate with potential confounders. a) Graph in Figure 3a with data points colored according to patient disease duration (longer duration: light blue; shorter duration: dark blue) (left), and plot showing correlation between disease duration and value on PC1 (right), as this component represents the largest difference between HC and MS. b-e) Same as (a) with MSSS (more severe score: light blue; less severe score: dark blue), age (older: light blue; younger: dark blue), sex (female: blue; male: red) and treatment status (HC: black; treated MS: grey; untreated MS: red) as potential confounding factors. f) Graph in Figure 3a with lines drawn between data points representing pairs of siblings. HC: healthy controls; MS: MS patients; MSSS: MS Severity Score.

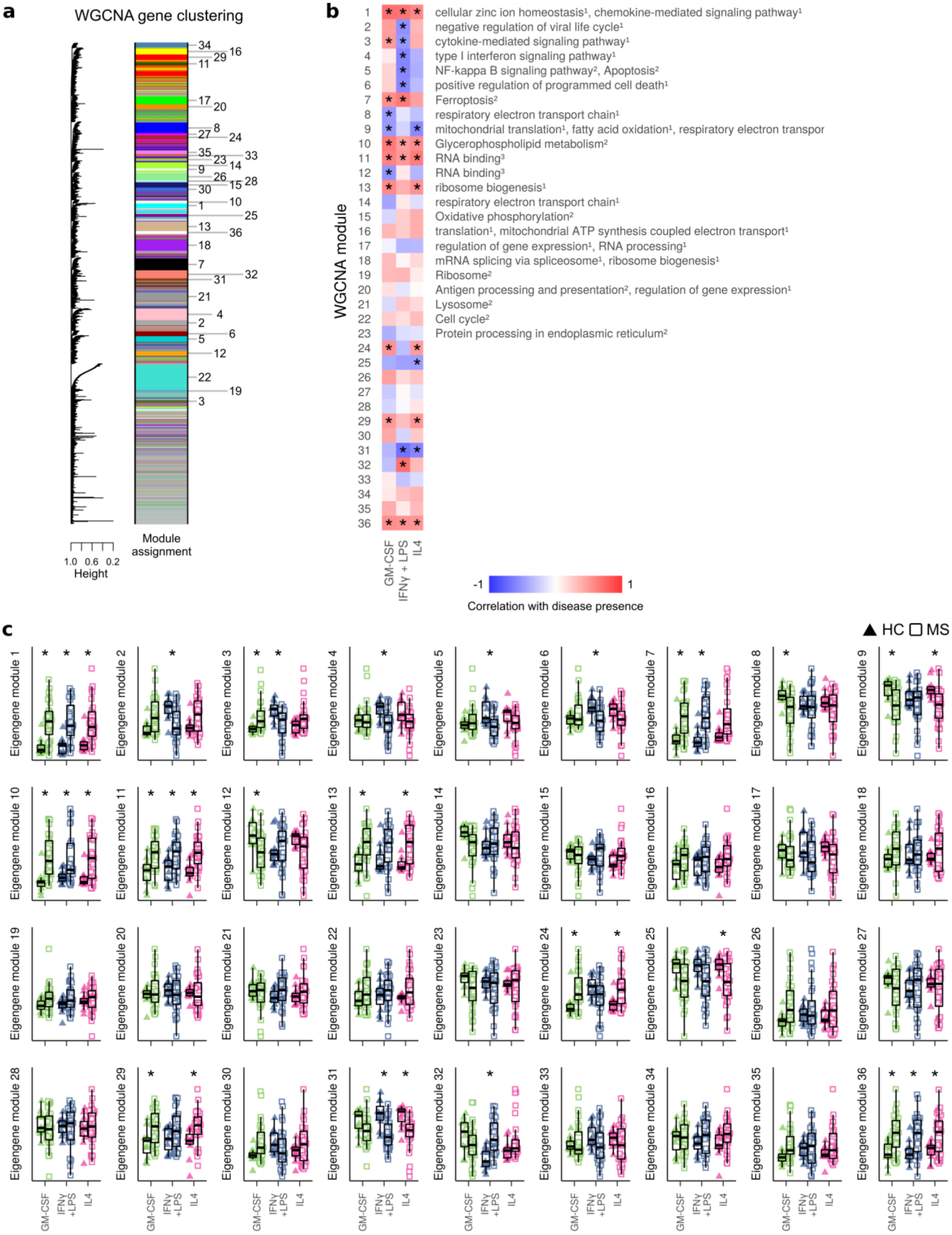
a
WGCNA gene clustering
Module assignment
1.0 0.6 0.2
Height
b
1 cellular zinc ion homeostasis¹, chemokine-mediated signaling pathway¹
2 negative regulation of viral life cycle¹
3 cytokine-mediated signaling pathway¹
4 type I interferon signaling pathway¹
5 NF-kappa B signaling pathway², Apoptosis²
6 positive regulation of programmed cell death¹
7 Ferroptosis²
8 respiratory electron transport chain¹
9 mitochondrial translation¹, fatty acid oxidation¹, respiratory electron transpor
10 Glycerophospholipid metabolism²
11 RNA binding³
12 RNA binding³
13 ribosome biogenesis¹
14 respiratory electron transport chain¹
15 Oxidative phosphorylation²
16 translation¹, mitochondrial ATP synthesis coupled electron transport¹
17 regulation of gene expression¹, RNA processing¹
18 mRNA splicing via spliceosome¹, ribosome biogenesis¹
19 Ribosome²
20 Antigen processing and presentation², regulation of gene expression¹
21 Lysosome²
22 Cell cycle²
23 Protein processing in endoplasmic reticulum²
WGCNA module
GM-CSF
IFNγ + LPS
IL4
-1
1
Correlation with disease presence
c
HC
MS
Eigengene module 1
Eigengene module 2
Eigengene module 3
Eigengene module 4
Eigengene module 5
Eigengene module 6
Eigengene module 7
Eigengene module 8
Eigengene module 9
Eigengene module 10
Eigengene module 11
Eigengene module 12
Eigengene module 13
Eigengene module 14
Eigengene module 15
Eigengene module 16
Eigengene module 17
Eigengene module 18
Eigengene module 19
Eigengene module 20
Eigengene module 21
Eigengene module 22
Eigengene module 23
Eigengene module 24
Eigengene module 25
Eigengene module 26
Eigengene module 27
Eigengene module 28
Eigengene module 29
Eigengene module 30
Eigengene module 31
Eigengene module 32
Eigengene module 33
Eigengene module 34
Eigengene module 35
Eigengene module 36
GM-CSF
IFNγ +LPS
IL4

**Fig. S5** Complete version of the WGCNA network shown in Figure 5. a) Clustering of genes and assignment of module labels. Module number is presented next to the largest clustering of module genes. b) Heat map of correlation between presence of disease and eigengenes of each module and activation. Functional annotations were given by Enrichr from databases GO biological process, GO molecular function, and KEGG pathways (adjusted $p<0.05$). c) Eigengene values in all modules, for each sample and module, grouped according to disease and activation state (HC: triangles; MS: squares; GM-CSF: green; IFNγ+LPS: blue; IL4: pink). HC: healthy controls; MS: MS patients (HC n=11; MS n=28). *$p<0.05$. Correlations tested with FDR-corrected default WGCNA functions.

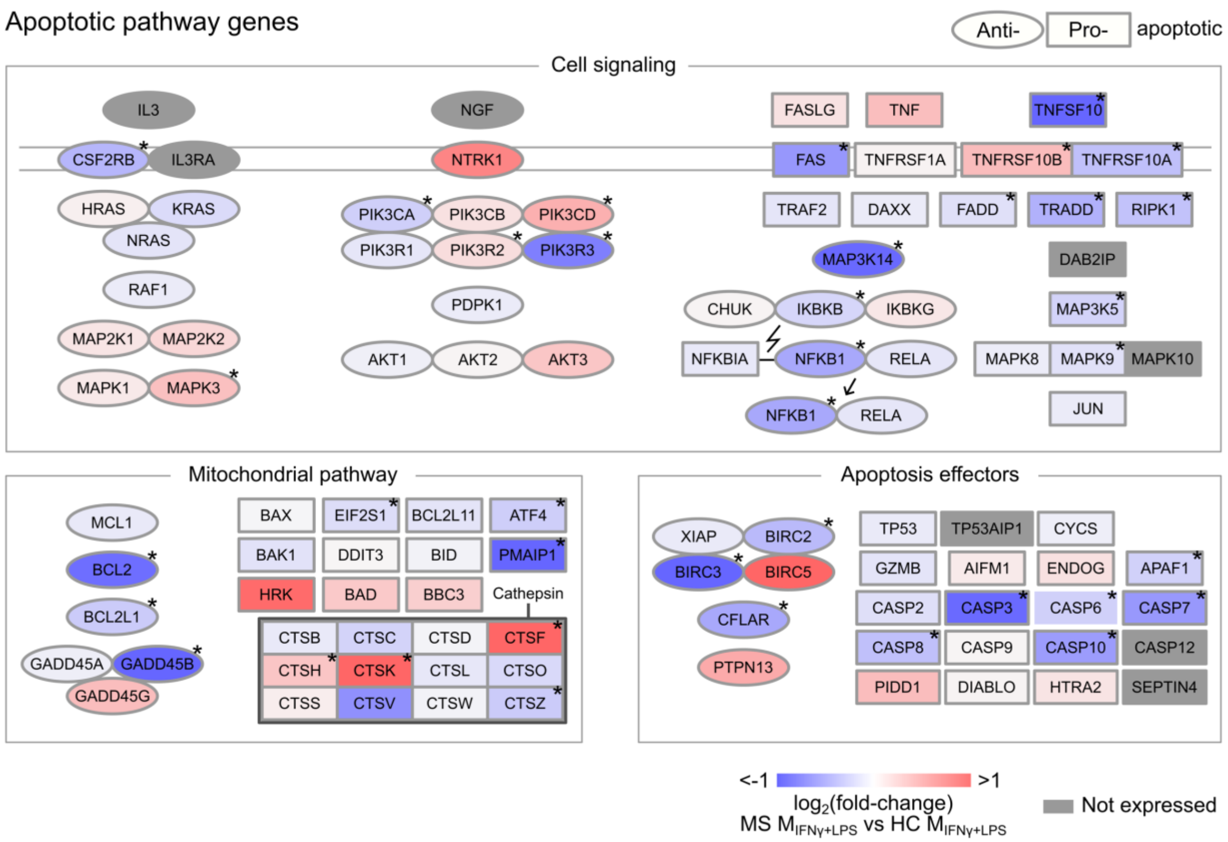


**Fig. S6** Illustration of all genes involved in signaling, mitochondrial response and effectors in the KEGG pathway Apoptosis. Each gene is colored according to the mean difference in expression between MS and HC $M_{IFN\gamma+LPS}$ (Δlog2(mean), over-expression in red, under-expression in blue, HC n=11; MS n=28). *significantly different genes (q<0.05) in limma differential expression analysis. HC: healthy controls; MS: MS patients.

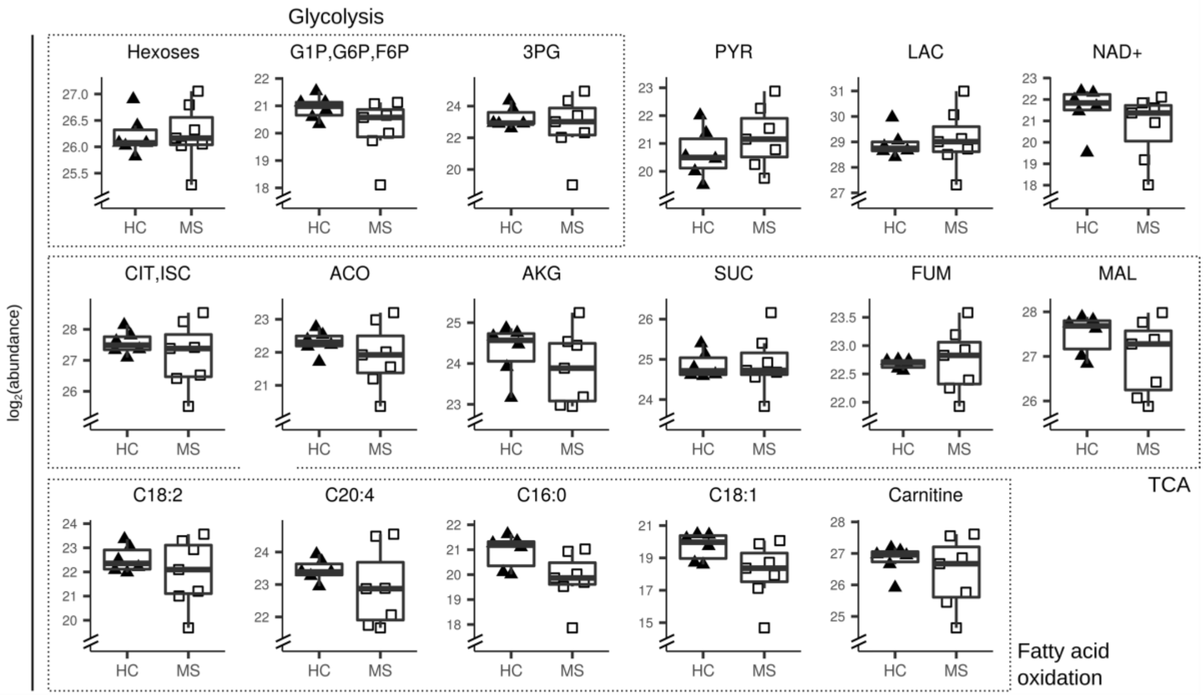


**Fig. S7** Boxplots of abundance of metabolites shown in Figure 6 in HC and MS. HC: healthy controls; MS: MS patients; G1P: glucose-1-phosphate; G6P: glucose-6-phosphate; 3PG: 3-phosphoglycerate; PYR: pyruvate; LAC: lactate; CIT: citrate; ISC: isocitrate; ACO: aconitate; AKG: alpha-keto-glutarate; SUC: succinate; FUM: fumarate; MAL: malate.
